# Cell type populations for 3D anatomical structures of the Human Reference Atlas

**DOI:** 10.1101/2025.08.14.670406

**Authors:** Andreas Bueckle, Bruce W. Herr, Lu Chen, Daniel Bolin, Danial Qaurooni, Michael Ginda, Yashvardhan Jain, Aleix Puig-Barbe, Kristin Ardlie, Fusheng Wang, Katy Börner

## Abstract

The human body contains ∼27-36 trillion cells of up to 10,000 cell types (CTs) within a volume of ∼62-120 liters (males) and 52-89 liters (females). The Human Reference Atlas (HRA) v2.3 provides a quantitative 3D framework of CTs across 73 reference organs and 1,283 3D anatomical structures (ASs). The HRA Cell Type Population (HRApop) effort has quantified CTs per AS using high-quality single-cell datasets processed through scalable, reproducible workflows and cell type annotation (CTann) tools. HRApop v1.0 includes reference CT populations for 73 ASs (112 when sex-specific) using 662 datasets spatially registered to 230 locations across 17 organs (31 when sex-specific). For 558 single-cell (sc-)transcriptomics datasets (11,042,750 cells), CTs and biomarker expressions were computed using Azimuth, CellTypist, and popV. To test generalizability, 104 sc-proteomics datasets (16,576,863 cells) were integrated. In total, HRApop includes 27,619,613 cells and serves as a healthy reference for researchers aiming to elucidate mechanisms underlying cellular interactions, FTU operations, and cellular and tissue level disease progression, which may facilitate advancements in basic discovery and lead to new therapeutic strategies.

## Background & Summary

### The Need

The volume^1^ of the adult human body is estimated to range from 62-120 liters (0.062-0.120 m³) in males with 36 or 37^2^ trillion cells to 52-89 liters (0.052-0.089 m³) in females with 27 trillion cells^3^. There is no consensus on the number of CTs within the human body. Estimates range from 400 major CTs^3–6^ to 3,358^7^ total CTs, and depend on the criteria used to determine what constitutes a CT (see the **Estimates of number of CTs in the human body** section for a more detailed discussion). Efforts like the Human BioMolecular Atlas Program (HuBMAP)^8,9^, the Human Cell Atlas (HCA)^10–12^, and many of the 20+ other atlas efforts contributing to the HRA endeavor aim to provide clarity based on high-quality experimental data collection and analysis.

Most atlas projects aim to capture the number and type of cells per AS together with biomarker expression values—based on expert knowledge or experimental data. For example, the Blue Brain Cell Atlas^13^ (bbp.epfl.ch/nexus/cell-atlas) features 3D data for the mouse brain with CT populations programmatically placed inside 737 brain regions defined in the Allen Mouse Brain Atlas^14^; different ASs; regions and their CT populations can be toggled on and off, and the color hue can be set to encode cell regions, types, or density; cell counts for neurons and glia (with confidence values) are also displayed. The Genotype-Tissue Expression (GTEx) sc-portal^15^ (gtexportal.org/home/singleCellOverviewPage) features CT populations from 25 tissue blocks in eight organs from 16 donors, and the Chan Zuckerberg Initiative (CZI) CELLxGENE (CxG) Portal^16^ (cellxgene.cziscience.com/datasets) features CT populations for 325 human, organ-level ASs but without visual representations of these structures. At the functional tissue unit (FTU) level^17^, Anatomograms^18,19^ developed by the Gene Expression Team at the European Molecular Biology Laboratory-European Bioinformatics Institute (EMBL-EBI) in collaboration with the Wellcome Sanger Institute; the Kidney Precision Medicine Project (KPMP)^20,21^ Explorer (atlas.kpmp.org/explorer); and the HRA FTU Explorer (apps.humanatlas.io/ftu-explorer) feature 2D medical illustrations with associated CT populations.

Given that life unfolds in 3D, there is a strong interest to capture CT populations for ASs in the human body in 3D. The HRA v2.3 features the 3D shape, size, location, and rotation of 1,283 3D ASs, which can be explored at humanatlas.io/3d-reference-library; each 3D AS belongs to one of 73 organs, also called 3D reference objects^22^. As new organ experts join the HRA effort, new 3D structures for male and female organs are added. Data from sc-portals such as GTEx, CxG, KPMP (atlas.kpmp.org), the HuBMAP Data Portal (portal.hubmapconsortium.org), or the Cellular Senescence Network (SenNet) Data Portal^23^ (data.sennetconsortium.org/search) can be spatially registered into the 3D reference objects using the web-deployed HRA Registration User Interface (RUI, apps.humanatlas.io/rui)^24,25^, guided by standard operating procedures (SOPs)^26–28^.

### The Challenge

Computation of CT populations for the many different ASs in the human body requires both a 3D registration and reproducible dissociation protocols that allow isolating single cells (or single nuclei) from tissue in support of sc-transcriptomics analyses^29–31^ or high quality cell segmentation for sc-proteomics spatial data^32–34^. In both cases, CTann tools are needed to assign a CT to each cell. Most sc-segmentation and annotation tools are organ specific. However, human organs are large (an average human kidney is about 10-12 cm long, 5-7 cm wide, and 3 cm thick and is estimated to contain about 110 billion cells^3^). Organs have many different internal ASs with vastly different CT populations that serve diverse physiological functions. The HRA^35,36^ used the Visible Human Project data^36–39^ and the expertise of medical illustrators to segment 1,283 3D ASs in 37 vital organs. Given tissue data that is spatially registered into these 3D AS using the RUI^24^ plus sc-dissociation/segmentations and annotations^32,40–48^, the number of cells per CT can be computed for specific ASs, i.e., 15 AS within the male lung. To compute HRApop at the AS (not organ) level, 16,293 datasets from four different portals were downloaded, donor metadata was harmonized, and multiple CTann tools were run for each sc-transcriptomics dataset. This paper explains how 662 high-quality datasets were chosen to compute dataset, extraction site, and AS-specific CT populations and biomarker expression values at scale, how open HRApop v1.0 data was published and what known limitations and next steps exist.

### The Opportunity

The 27,619,613 cells in HRApop v1.0 represent CT populations for 73 ASs (112 when sex-specific) across 17 organs (31 when sex-specific). CTs and biomarker expressions were computed using Azimuth^49^, CellTypist^50,51^, and/or popV^52^ for sc-transcriptomics datasets. These can serve as an AS-level, healthy reference for stakeholders working on the processes behind cell-to-cell communication, FTU activity, and disease advancement in cells and tissues, which could enable progress in foundational research and the development of innovative therapies. To make the HRApop v1.0 dataset maximally available for single-cell biologists, bioinformaticians, computational biologists, and physician scientists, it was made available as 5-Star Linked Open Data (LOD)^53^, with full provenance including donor metadata and resolvable Uniform Resource Identifier (URI), making it a Findable, Accessible, Interoperable, and Reusable (FAIR)^54^ resource.

In addition to downloading data products and downstream usage in various HRA applications and the HRA Application Programming (API, apps.humanatlas.io/api), SPARQL (www.w3.org/TR/sparql11-query/) queries can be run, e.g., to retrieve all AS-CT combinations, with sex, tool, CT, and cell percentage as CSV files (see **Data Records** section). Beyond direct download, the **Usage Notes** section points to Python example code for using the HRA API to get <u>A</u>natomical <u>S</u>tructure Cell Type <u>Pop</u>ulations (ASpop) as well as <u>D</u>ataset and <u>E</u>xtraction <u>S</u>ite Cell Type <u>Pop</u>ulations (DESpop) via the HRA KG^22^, see **Box 1**. A companion website for this paper is available cns-iu.github.io/hra-cell-type-populations-supporting-information.

## Overview

The HRApop effort combined (1) CT populations from sc-transcriptomics and sc-proteomics data made with CTann tools (or via user-assigned CT label), (2) donor metadata, and (3) 3D extraction sites made via the RUI. This involved running two scalable workflows, optimized to efficiently deal with the continuously growing number of datasets. Over the last three years, the processes used to <u>D</u>ownload data, perform <u>C</u>ell <u>T</u>ype <u>A</u>nnotations, and compute CT populations per dataset (called **DCTA Workflow**) and the workflow that takes CT populations, donor metadata, and 3D extraction sites via the <u>RUI</u>, <u>to</u> compile <u>C</u>ell <u>T</u>ype <u>Pop</u>ulations for ASs, datasets, and extraction sites (called **RUI2CTpop Workflow**) were implemented. Note that CT populations recorded the number of cells per CT not just for ASs but also for datasets and extraction sites as well as the top-10 biomarkers per CT (datasets only) and their mean expression values (see **Methods** section on how these were computed).

Subsequently, this **Background & Summary** section details a generalization to sc-proteomics data, then concludes with limitations and outlook for future work on improving the HRApop Atlas Data presented. In the remainder of the paper, the **Methods** section details relevant terminology, describes implementation of the DCTA and RUI2CTpop Workflows, and the used datasets and their sources are described. The **Technical Validation** section shows confidence scores per cell per tool, gene counts, the prevalence of different CTs inside not just the organs but also the ASs of the human body, and the number of datasets per organ and AS.

## Generalization to spatial data

In HRApop v1.0, 104 sc-proteomics datasets contributed 16,576,863 cells, which brought the total number of cells to 27,619,613. They were associated with high-quality publications^35,55–59^ and used protein and antibody-based modalities such as cyclic Immunofluorescence (CycIF)^60^, Cell DIVE^61–63^, and co-detection by indexing (CODEX)^64^ to identify proteins and quantify their expressions in a tissue in situ; there are plans to integrate iterative bleaching extends multiplexity (IBEX)^65,66^ datasets in the future. While this paper does not focus on sc-proteomics data, it presents a curated collection of sc-proteomics datasets as a generalized use case for HRApop. Spatial proteomics has received significant interest from the scientific community in recent years^67^, which has led to increased high-quality data generation. If integrated in HRApop, it enables creating CT populations with a preserved spatial context for each cell —which is lost in sc-transcriptomics datasets, although some recent work has attempted recoveries for specific assay types and organs^48^.

The DCTA Workflow has output CT populations and metadata for all datasets, both sc-transcriptomics and sc-proteomics. While sc-transcriptomics datasets were run through at least one CTann tool, sc-proteomics datasets had CTs assigned by their expert providers, who shared CT populations in the form of CSV files, see code in **Table S1**. The DCTA Workflow systematically went through a list of sc-proteomics datasets (available on GitHub^68^) and transformed the files with nodes for each cell in each dataset (CSV) into a CT population (JSON) as an input for the RUI2CTpop Workflow (see GitHub^69^). In the future, the DCTA Workflow will be extended to (1) handle other generalized use cases that cannot be annotated with CTann tools and (2) run more CTann tools over new and existing HRApop datasets for additional CT populations.

To make crosswalks for sc-proteomics data, CT labels were shared by contributors of high-quality datasets and documented in a related publication^34^; crosswalks for each dataset were manually created for that publication effort and made available at lod.humanatlas.io/ctann/vccf/latest. Unmapped CTs are on GitHub^70^.

## Estimates of number of CTs in the human body

While no consensus exists on the number of CTs within the human body, estimates range from 400 major CTs^3–6^ to 3,358 CTs^7^, depending on the criteria used to determine what constitutes a CT. As an example, in the retina, a major class of retinal neurons is the amacrine cell (purl.obolibrary.org/obo/CL_0000561). This cell can be subdivided in multiple ways. At a broad level, it is often classified into GABAergic (purl.obolibrary.org/obo/CL_4030027) and glycinergic (purl.obolibrary.org/obo/CL_4030028) types (two categories). However, based on morphology, researchers have identified 25 distinct amacrine CTs^71^. At the level of sc-transcriptomics, the Human Retina Cell Atlas (HRCA)^72^ has reported 123 CTs in the retina, including 73 molecularly distinct amacrine CTs. As a result, depending on the resolution—functional class, morphology, or transcriptomic profile—one might count one, two, 25, or 73 different amacrine CTs. In the case of the human brain, sampling more than three million nuclei from approximately 100 dissections across the forebrain, midbrain, and hindbrain, 461 clusters and 3,313 subclusters (granular CTs) organized largely according to developmental origins were identified^73^. Combining CTs in the Human Lung Cell Atlas^74^ and the CellRef atlas from the LungMAP consortium^75^ revealed 68 distinct CTs in the human lung and nasal cavity^76^. As of August 2025, CL^77^ contains 3,358 classes, but this includes many non-human CT terms as well as grouping classes—i.e., internal nodes in the hierarchy that do not correspond to distinct, terminal CTs. When limiting the count to leaf-level human CTs, the number is on the order of 2,500, but this does not reflect many of the novel CTs that have been recently defined using single-cell technologies. Further, about 2,000 CTs^7^ in Cell Ontology (CL, www.ebi.ac.uk/ols4/ontologies/cl)^77–79^ are connected to ASs in the cross-species anatomy ontology Uberon^80,81^ via ‘part of’ relationships. As a result, 400 total CTs might be a clear underestimation. Assuming that there are 78 major organs in the human body of varying size and cellular complexity, with most organs averaging between 50 and 120 transcriptomic CTs, aside from the brain with 3,000-5,000, there might be close to 10,000 CTs in adult mammalian organisms, depending on the criteria for distinguishing CTs.

## Limitations

HRApop v1.0 comes with a number of limitations:

### Dataset duplication across data portals

Some datasets (e.g., by KPMP) are available via multiple data portals (e.g., HuBMAP and CxG). However, no dataset should be used twice for HRApop construction. Data duplication detection is difficult as different versions of the data might exist on different portals and different metadata might be associated with each version. For example, there might be a dataset submitted with the very first paper submission that is linked to a preprint, a slightly expanded dataset associated with a revised and later preprint version, and a final version of the dataset linked to a peer-reviewed published paper; paper title and authors might also change in the process making deduplication challenging.

### CTann tools were trained on underspecified data

Azimuth, CellTypist, and popV were trained on high-quality reference datasets that might not have been separated by sex or other donor demographics, and for which RUI extraction sites were not available; i.e., it is unknown from which of the diverse ASs within an organ a tissue that was used to train the CTann tool originated. HRApop, however, utilizes existing CTann tools to compile ASpop that are specific to the diverse ASs and tailored to the male and female human body. As more RUI registered sc-transcriptomics datasets become available, CTann tool developers might like to use this additional tissue origin information to optimize CTann training and CT annotation predictions.

### Missing assay type information needed for batch correction

At present, the ds-graph HRA Digital Object type^22^, which captures datasets and their extraction sites plus donor metadata, only contains assay type metadata as free text string rather than not ontology terms (e.g., 0x 3’ v1-3, 10x scATAC-seq, MERFISH, Smart-seq, all listed on the CxG portal). The HRA KG will be extended to provide look-up tables for assay types to ontology terms for enabling more systematic queries. Also, collaboration is ongoing with the HuBMAP and SenNet portal teams to utilize ontology terms for identifying and standardizing assay types.

### Intersecting 3D reference objects

The 1,283 3D ASs are supposed to have no overlap with each so that CT populations are specific to one, not multiple intersecting ASs. However, in HRA v2.3, 18 ASs in 17 organs (e.g., two in the female heart) have intersections with each other and also have extraction sites in them (labeled “TB3” in **Fig. S1**). The AS-AS pairs with the most (here three) extraction sites that collide with both ASs are the ‘outer cortex of kidney’ and the ‘renal pyramid A’ in the male, left kidney and the ‘kidney capsule’ and the ‘outer cortex of kidney’ in the female, left kidney. Intersections will be corrected and tissue blocks will be re-registered for an upcoming HRA release.

### Limited coverage of organs and Ass

HRApop aims to enrich all 1,283 3D ASs in 73 3D reference objects of the entire HRA v2.3 with CT populations from high-quality datasets. However, in v1.0, only 73 3D ASs (112 if male and female are counted separately) in 17 organs (31 if male and female are counted separately) had data to compute AS-specific CT populations. The **Technical Validation** section expands on current HRApop coverage of organs and ASs.

## Outlook

To improve coverage and quality of HRApop, the following steps are planned:

### Long-term sustainability

The HRA and, by extension, HRApop is currently funded via the HuBMAP, SenNet, KPMP, Common Fund Data Ecosystem (CFDE), GTEx, and NIDDK. Funding that has been acquired since HRApop v1.0 was released comes via the Whole Person Reference Physiome Research and Coordination Center (WPP, [1U24AT013504-01](https://reporter.nih.gov/search/TJAFVzPxvk64YuUPEy5Rlw/project-details/11224772)), the Canadian Institute for Advanced Research (CIFAR) MacMillan Multiscale Human (cifar.ca/research-programs/cifar-macmillan-multiscale-human), and the Stiftung Charité via Berlin Institute of Health at Charité (BIH). This funding supports regular HRApop releases, with data products made available via Zenodo^82^ and the HRA KG^22^ (lod.humanatlas.io/graph/hra-pop).

### Increase number of RUI registered datasets

As of December 2025, the HuBMAP and SenNet portals list 799 and 272 datasets that are currently in QA/QC status but will be published soon. Outreach efforts to authors of peer-reviewed, published papers are ongoing to register their data for use in the DCTA Workflow. The HRApop effort will also integrate data from Tabula Sapiens^5^, KPMP, the Helmsley Gut Cell Atlas^83^, and the Deeply Integrated human Single-Cell Omics (DISCO) database^84^, which has a total of 21,330 datasets, out of which 32.8% of the total data is from a healthy human body, across 166 unique ASs. Other potential sources for high-quality datasets have been captured in Hemberg et al.’s recent article on large cell atlases^85^.

### Scale up tissue registration via millitomes

A millitome^35^ (from Latin *mille*, meaning “thousand,” and the Greek *temnein*, meaning “to cut”) is a device designed to hold a freshly procured organ and facilitate cutting it into many small tissue blocks of well defined size for usage in sc-analysis and HRA construction. It is used to produce uniformly sized slices or cubes of tissue material that can be registered to 3D reference objects. Using a millitome improves efficiency by enabling consistent, high-throughput sampling. Recently, 209 brain tissue blocks have been added and will be included in the next HRApop run as part of the 10th HRA release.

### Improve generalizability

The 104 sc-proteomics datasets in HRApop v1.0 were presented as a generalization from sc-transcriptomics datasets. In the future, and in synergy with HRA Vasculature Common Coordinate Framework (VCCF) construction efforts^57,86^ around endothelial cell environments^34^, more CODEX^64^ datasets and new modalities, such as the Spatial Multiomics Single-Cell Imaging platform CosMx^87^, will be added to HRApop. A recent paper^34^ analyzed 399 spatially resolved omics datasets from 14 studies comprising 12 tissue types with a total of 47,349,496.

### Increase number of CTann tools used to enable more benchmarking

Currently, HRApop uses three well-established CTann tools backed by scientific publications describing the methods, results, and validations for each tool. Results are presented as CT populations by CTann tool—users can pick their favorite tool and data or perform comparisons and benchmarks between CTann tools^76^. Future HRA releases will feature additional CTann tools such as FR-Match^88^ and Pan-Human Azimuth (satijalab.org/pan_human_azimuth) in support of improved cell type annotation, CTann tool comparisons and benchmarking.

### Add biomarker sets

For HRApop v1.0, the top biomarkers per CT per dataset were computed using *scanpy*’s *rank_gene_groups()* method (scanpy.readthedocs.io/en/stable/generated/scanpy.tl.rank_genes_groups.html). In future HRApop releases, additional sets of top biomarkers will be provided to the user by running, e.g., NS-Forest^89^ during the annotation phase of the DCTA.

### Decrease run time for HRApop code

For HRApop v1.0, the DCTA Workflow started on Thu, May 15, 2025, ran for about 10 days, and finished on Sunday, May 25, 2025. It averaged 87.63 dataset annotation runs per hour. Annotations took about 8.59 days to finish. This long runtime is primarily due to the complexity of the annotation pipeline, including annotation, crosswalking, and mean gene expression per cell type, to cover over 10,000 datasets and tens of millions of cells. Targeted optimization of the algorithms and workflows combined with more hardware resources and re-using annotations from prior runs will be required to reduce runtime. Work is underway to save annotations between runs to skip the re-annotation step. After a 22-day QA phase, the RUI2CTpop Workflow started on June 16 at 5:55:27 PM EDT and finished about four hours later at 10:07:11 PM EDT the same day. A full log is linked in **Table S1**. In the future, the run time for the DCTA Workflow will be decreased by using high performance computing (HPC), e.g., Big Red at Indiana University (kb.iu.edu/d/brcc). Also, the crosswalking will be moved to the RUI2CTpop Workflow, which will decrease runtime and increase the modularity of both workflows.

## Methods

### Terminology

**Box 1** introduces key terminology for constructing and using HRApop; it also provides links to concepts already defined in related papers.

#### Box 1.

Terms in *italics* have their own entry, either in this Box or in the HRA marker paper^35^ or the HRA KG paper^22^. The former already contains definitions for: *3D collision, extraction site, AS, ASCT+B table, biomarker, CT, CTann tool, crosswalk, millitome, polygon mesh, registration set, RUI registered, sc-transcriptomics/sc-proteomics data, tissue block,* and *(weighted) cosine similarity*. The latter contains definitions for: *3D reference object, ds-graph*, and *SPARQL (*www.w3.org/TR/sparql11-query).

- **<u>A</u>natomical <u>S</u>tructure Cell Type <u>Pop</u>ulations (ASpop)**: Captures the number of cells per CT for an *AS* given with an ontology ID, donor sex, *AS* label, the *CTann tool* used to assign *CTs*, and a list of datasets from which the data was sourced. An exemplary *ASpop* is shown on the companion website at cns-iu.github.io/hra-cell-type-populations-supporting-information/#for-an-as.
- **Cell instance**: Is an occurrence of a unique cell, identified by its bar code or a similar unique ID. In the data products of HRApop v1.0 (see **Data Records**), cells can be annotated by multiple CTann, which leads to potentially multiple occurrences of the same cell.
- **Cell type population**: Is a listing of unique CTs and their counts computed for *ASs, extraction sites*, and datasets. For the latter, mean B expression values are also computed. *CT populations* are computed from *CT* counts in experimental datasets, obtained either via *CTann* in the *DCTA Workflow* (for *sc-transcriptomics* datasets), or via expert/author-provided annotations (*sc-proteomics* datasets). *CT populations* are associated with exactly one *CTann* tool; if a dataset has *CT populations* from multiple *CTann* tools, the *CTann* tool that computed the CT population becomes part of its provenance. *CT populations* are kept inside the *ASpop* graph (for an entire AS) and the *DESpop* (for datasets and extraction sites).
- **Collision detection (bounding box)**: Is a computationally efficient but imprecise method for identifying intersections between two 3D objects in Euclidean space. For each 3D object, the bounding box is a cuboid shape that is calculated that encapsulates all vertices and only this bounding box is used when computing collisions.
- **Collision detection (mesh-based)**: A more precise but computationally more expensive method for identifying the intersection between two 3D objects in Euclidean space based on mesh surfaces, resulting in *AS* tags for extraction sites. To enable mesh-based collision detection, *3D reference objects* need to be pre-processed. Details are provided in the **Methods** section.
- **Corridor**: Describes a 3D volume that encompasses all possible locations for an *extraction site* while maintaining its *intersection percentages* with any *ASs* based on *mesh collision detection*.
- **Criteria C1-4**: Describes four conditions that a dataset must fulfill in order to be used for HRApop construction: (1) it has an extraction site in a 3D reference object, (2) it has a *CT population* via *CTann* or the *sc-proteomics* workflow, (3) it comes from a data portal with QA/QC or have an associated peer-reviewed publication in a scientific journal, and (4) it is from a healthy adult human donor.
- **<u>D</u>ataset and <u>E</u>xtraction <u>S</u>ite Cell Type <u>Pop</u>ulations (DESpop, formerly “Atlas-Enriched Dataset Graph” in a related HRA publication**^35^): Is a collection of datasets that meet *Criteria C1-4* and are used for HRApop construction. A snippet of the *DESpop* is shown on the companion website at cns-iu.github.io/hra-cell-type-populations-supporting-information/#for-a-dataset.
- **<u>D</u>ownload and <u>C</u>ell <u>T</u>ype <u>A</u>nnotation (DCTA) Workflow**: Is a set of scripts to download H5AD files, execute the Docker (www.docker.com) containers in *HRApop CTann Tool Containers*, and output CT populations and data metadata as input for the RUI2CTpop Workflow.
- **Donor**: Is an individual person, living or deceased, who contributed tissue. Donors have demographic metadata, such as age, body mass index (BMI), race/ethnicity, and sex. All *donors* in HRApop are healthy, human adults.
- **Experimental dataset**: Describes cell by gene matrices (via *CTann tools*) or cell by protein matrices via Organ Mapping Antibody Panels (OMAPs)^90^ derived from an organ-specific *tissue block* from a *donor*.
- **HRApop Atlas Data (also called HRApop Data Used in Atlas Construction)**: Is a collection of high-quality experimental datasets that fulfill *Criteria C1-4* that is used in atlas construction. Also part of the *HRApop Atlas* are the *CT populations of* 3D extraction sites of these datasets, plus the ASs with which these extraction sites collide. At the beginning is a programmatically compiled collection of dataset IDs, extraction sites, and donor metadata that forms the input of the *DCTA Workflow*. Dataset IDs can come from three sources: (1) sc-portals with access to sc-transcriptomics H5AD files (from HuBMAP, SenNet, CxG, and GTEx), which are then given to the *DCTA Workflow* as an input, (2) sc-proteomics datasets, and (3) HRA Registrations, see GitHub^91^ and **Table S1**, a set of manually curated extraction sites, from which the *RUI2CTpop Workflow* extracts dataset IDs and attempts to match them with *CT populations* from the *DCTA Workflow*. Each HRApop run provides a series of reports (see GitHub^92^).
- **HRApop CTann Tool Containers**: Is a collection of CTann tools containerized with Docker to annotate H5AD files. The DCTA Workflow uses the *HRApop CTann Tool Containers*. They also contain utilities for CTann, e.g., computing gene expressions and identifying the top-n genes.
- **Intersection percentage or volume**: Is a measure that describes the shared 3D space between two 3D objects (e.g., a *tissue block* and an *AS*) expressed as a percentage of the total volume or as an absolute value in cubic millimeters (mm^3^).
- **Mean biomarker expressions**: Are computed for the top 10 genes per CT per dataset, then stored in the *DESpop*.
- **<u>RUI</u> to Compile <u>CTpop</u> (RUI2CTpop) Workflow**: Is a collection of scripts to compile the *HRApop Atlas Data* from the output of the *DCTA Workflow* (see **Fig. 1 and Methods**).

**Fig. 1.**
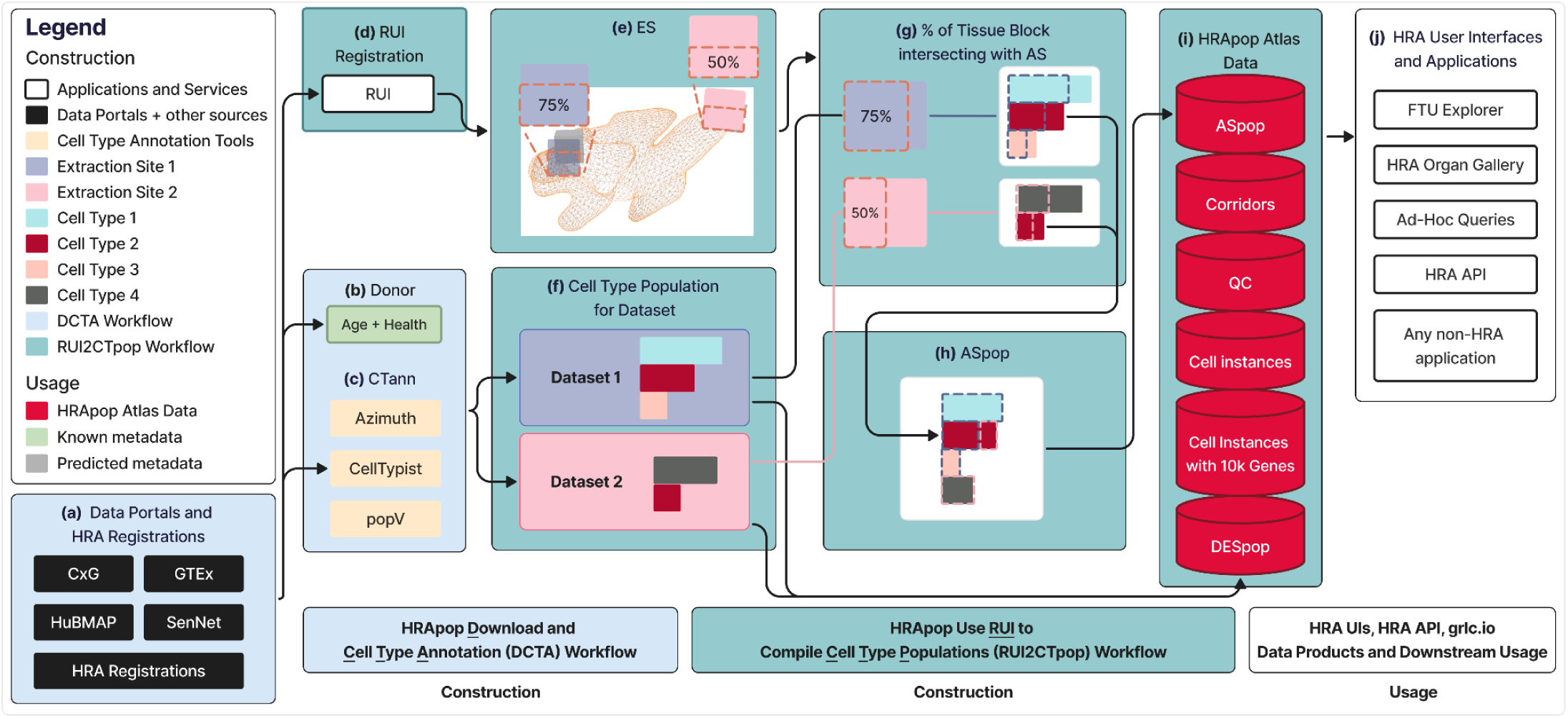
Construction and usage of HRApop. **(a)** Data is ingested from four data portals or from HRA Registrations^91^. **(b)** Donor information is extracted from dataset metadata or the extraction site (if from HRA Registrations). **(d)** CT populations are computed using CTann tools if possible. **(a)-(c)** is handled by the **DCTA Workflow**. **(d)** RUI-assigned extraction sites for each dataset are identified. If a dataset has an extraction site, **(e)** intersection percentages (see **Box 1**) of the extraction site with ASs can be computed. **(f)** CT populations for corresponding datasets obtained via one or more CTann tools can be combined with **(g)** intersection percentages between extraction site and AS, resulting in **(h)** ASpop, which, together with DESpop, are then published as **(i)** HRApop Atlas Data. **(d)-(i)** is handled by the **RUI2CTpop Workflow**. **(j)** illustrates usage of HRApop Atlas Data by applications inside and outside the HRA data ecosystem.

## Construction

**HRApop** was constructed with two automated workflows:

1. The **<u>D</u>ownload and <u>C</u>ell <u>T</u>ype <u>A</u>nnotation (DCTA) Workflow** (see GitHub^93^ and **Box 1**) used scalable, open-source software to programmatically download and annotate H5AD files from four portals: HuBMAP, SenNet, GTEx, and CxG. Cells in these H5AD files were then annotated with containerized CTann tools kept in the **HRApop CTann Tool Containers** (see **GitHub**^94^ and **Box 1**); next, the DCTA Workflow crosswalked the resulting CTann labels to CL^77^ or Provisional Cell Ontology (PCL)^95^, see **Crosswalks** section for details, then compiled donor metadata from each dataset and finally made the result available as a set of CSV files (for metadata) and JSON files (for CT populations). These could then be used as input for the next workflow.
2. The **<u>RUI</u> to Compile <u>CTpop</u> (RUI2CTpop) Workflow** (see GitHub^96^ and **Box 1**) then computed CT populations for ASs, datasets, and extraction sites. It first identified all datasets that could be used to construct HRApop based on four Criteria (C1-4, see **Box 1**):
  ○ **C1:** The dataset had a 3D extraction site by registration with the RUI (see **Methods**), i.e., the spatial position, rotation, and size of the tissue block (from which the dataset had been derived) within the HRA reference system is known and AS tags exist.
  ○ **C2:** The dataset had a CT population (see **Box 1**), i.e., number of cells per CT. For sc-transcriptomics data, this meant having an associated H5AD file with a cell by gene matrix that could be annotated via Azimuth^49^, CellTypist^50,51^, and/or popV^52^; for sc-proteomics data, CT populations were obtained via manual segmentation and annotation workflows.
  ○ **C3:** The dataset was of high quality, i.e., it had been downloaded from a data portal with built-in quality assurance/quality control (QA/QC) or was associated with a peer-reviewed publication.
  ○ **C4:** The dataset had complete donor metadata, was from a healthy tissue sample, and had an age value greater than 18. That is, the data was from a healthy adult human.

**C1** and **C2** were enforced to enable usage of the dataset to fill 3D AS with cells from one or multiple CTann tools. **C3** ensured that atlas data came from reliable sources. **C4** was in support of constructing a healthy reference for HRA User Story (US) #3 (compare reference tissue with aging/diseased tissue)^35^.

For datasets that met **C1-4**, RUI2CTpop output the **HRApop Atlas Data** (also called “**HRApop Data Used in Atlas Construction**,” see **Box 1**), a collection of data products to describe high-quality ASpop and DESpop for the HRA. Datasets that did not meet all four criteria were disregarded.

**Fig. 1** shows construction and usage of HRApop, from the data download on the left to the publication and usage of HRApop Atlas Data on the right.

First, in the **DCTA Workflow** (blue), datasets represented as H5AD files were programmatically downloaded from the four data portals (**Fig. 1a)** alongside donor metadata (**Fig. 1b**). Before download, non-human and diseased data were filtered out. Then, each dataset was annotated using all applicable CTann tools (**Fig. 1c**), resulting in CT populations if the dataset originated from a supported organ (**Fig. 1f**). Only if a dataset met all Criteria C1-4 (see **Box 1**), then it was used for HRApop Atlas construction.

Next, in the **RUI2CTpop Workflow** (green), datasets were matched against all existing RUI extraction sites by their ID (**Fig. 1d)**; the extraction sites held metadata on organ sex and laterality (left or right) and could come from either an API (such as for HuBMAP or SenNet, see **HRA registrations and extraction sites via APIs** section) or the static HRA Registrations^91^ (see **Table S1**), a manually curated collection of extraction sites.

For each HRApop Atlas Dataset, i.e., each that fulfilled Criteria C1-4 (see **Box 1**), the 3D ASs that its extraction site collided with were determined via mesh-based collision detection (**Fig. 1e**, see also **Box 1)**. The example shown is the renal pyramid (purl.obolibrary.org/obo/UBERON_0004200) of the left, male kidney (lod.humanatlas.io/ref-organ/kidney-male-left/latest) with two hypothetical tissue blocks colliding with it at 75% and 50% of the total volume of the extraction site. The number and percentage of CTs that should be in the colliding AS based on the intersection percentage (see **Box 1**) of the extraction site were computed (see **Fig. 1g**). Typically, many tissue blocks from different donor demographics (e.g., age, ethnicity) existed per male/female-specific ASs. Often, research teams carefully sample from the very same RUI extraction site (see definition in a related publication^35^) so hundreds of datasets could have the very same extraction site. This was performed for every extraction site that intersected with the AS. The result is an ASpop containing the unique and shared CTs contributed by each colliding extraction site (see **Fig. 1h**).

Finally, the ASpop and the DESpop used to generate the ASpop were published as two separate Resource Description Framework (RDF, www.w3.org/RDF) graphs (see **Fig. 1i**) via the HRA KG^22^ at lod.humanatlas.io/graph/hra-pop/latest in support of LOD principles.

Finally, the data products generated as part of this RUI2CTpop Workflow were made available for usage in various HRA UIs and the HRA API (apps.humanatlas.io/api, **Fig. 1j**). HRA KG queries can be run to support diverse applications inside and outside the HRA data ecosystem. Examples are the HRA FTU Explorer (apps.humanatlas.io/ftu-explorer), the HRA Organ Gallery in virtual reality (VR)^97,98^, and ad-hoc queries, such as one that provides an overview of all AS-CT combinations, including sex, tool, cell count, and cell percentage, available at apps.humanatlas.io/api/grlc/hra-pop.html#get-/cell_types_in_anatomical_structurescts_per_as.

An exemplary ASpop from the kidney and an exemplary snippet of the DESpop are available on the companion website at cns-iu.github.io/hra-cell-type-populations-supporting-information#exemplary-cell-summaries. A complete listing of all data and code for construction and usage of HRApop is provided in **Table S1.** An overview of all HRA applications that use **HRApop Atlas Data** is provided in **Table S2.**

## HRA registrations and extraction sites via APIs

For all extraction sites, mesh-based collision detection (see **Box 1**) was used to compute the intersection percentage (see **Box 1**) with 3D ASs, which enabled the construction and aggregation of CT populations for these 3D ASs. As a result, extraction sites and ASs in the HRA were enriched with the number of cells per CT from high-quality data. Where possible, for all datasets, extraction sites, and ASs, HRApop provided CT populations from every tool that could annotate the dataset. All extraction sites used in the HRApop Atlas Data are shown in the HRApop-focused Exploration User Interface^24^ (EUI) at cns-iu.github.io/hra-cell-type-populations-supporting-information/eui.html. By working closely with authors of published, high-quality datasets, as well as tissue providers in HuBMAP and SenNet, RUI was used to generate extraction sites for a growing number of organs and ASs.

**Table 1** presents counts for datasets, extraction sites, ASs, and organs based on input data for RUI2CTpop Workflow and the HRApop Atlas.

**Table 1.**
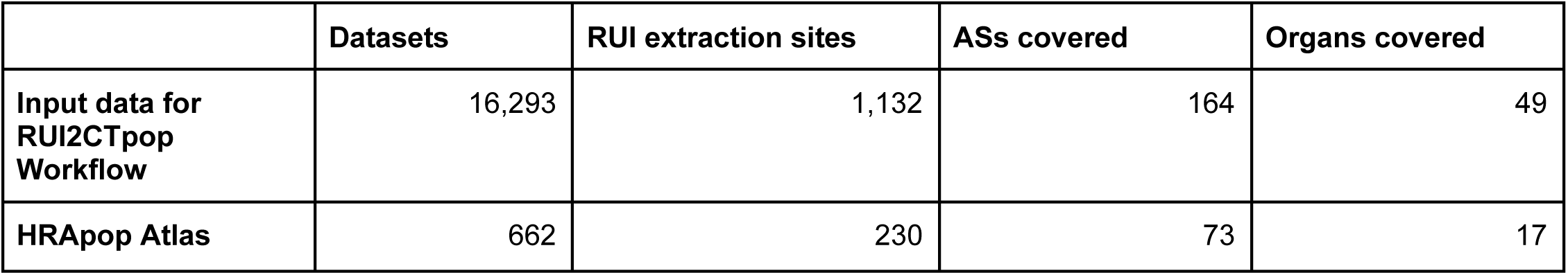
Number of datasets, extraction sites, ASs, and organs covered in HRApop v1.0.

|  | Datasets | RUI extraction sites | ASs covered | Organs covered |
| --- | --- | --- | --- | --- |
| Input data for RUI2CTpop Workflow | 16,293 | 1,132 | 164 | 49 |
| HRApop Atlas | 662 | 230 | 73 | 17 |

## Experimental data

The Sankey diagram in **Fig. 2** provides a high-level overview of the input data for the input for RUI2CTpop Workflow along several axes. It can be explored interactively at cns-iu.github.io/hra-cell-type-populations-supporting-information/sankey_universe_plotly.html. A version showing only HRApop Atlas Data is shown in **Fig. S2**, with an interactive version available at cns-iu.github.io/hra-cell-type-populations-supporting-information/sankey_atlas_plotly.html.

**Fig. 2.**
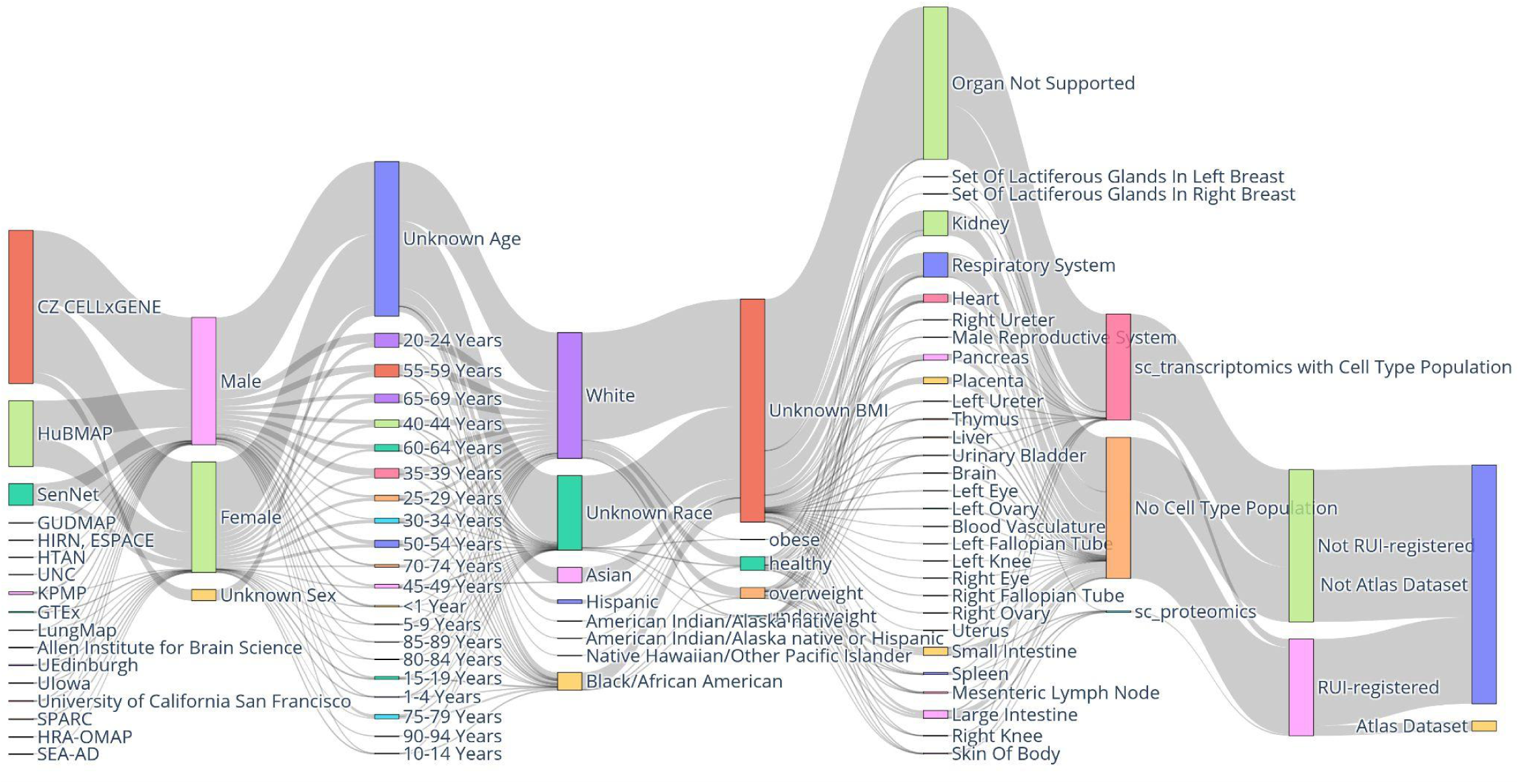
Sankey diagram of all input data for the RUI2CTpop Workflow. Note the Atlas Data node in the bottom right corner, which comprises the 662 HRApop Atlas Datasets in HRApop v1.0.

The Sankey diagram has 9 vertical axes that represent:

### Portal/source

Identifies the effort where the data originated. H5AD files were downloaded from HuBMAP, SenNet, GTEx, and CxG. The majority of datasets came from CxG. All other portals/sources were derived from extraction sites.

### Donor sex, donor age, donor race, donor body mass index (BMI)

Describe clinical metadata for the human specimens from which the data was retrieved. Where available, donor age was provided as an integer by HuBMAP and SenNet but as string by CxG (e.g., “61-year-old human stage”); as a result, string literal age values for CxG data were parsed as a number where possible. To enable visualization in a Sankey diagram, all age values were aggregated into bins of five years. For race, the same categories were used as on the HuBMAP Data Portal (portal.hubmapconsortium.org) as of the writing of this paper. BMI values were mapped to categories by brackets defined by the Centers for Disease Control on www.cdc.gov/bmi/adult-calculator/bmi-categories.html.

### Organ

Indicates the organ of origin. “Organ Not Supported” means that there was no matching 3D reference object in the HRA (e.g., for blood).

### CT population

Means that a dataset had either (1) one or multiple CT populations from one or multiple CTann tools (“sc_transcriptomics with Cell Type Population”), (2) no CT population because no CTann tool existed for the dataset (“No Cell Type Population”), or (3) a CT population via sc-proteomics as a generalization of the HRApop workflow (“sc_proteomics with Cell Type Population”).

### Extraction site

Indicates whether a dataset had an extraction site via RUI registration or not.

### HRApop Atlas Data

Indicates whether the dataset was part of the HRApop Atlas or not This was only true for 662 datasets.

## Computational resource requirements

For both the DCTA and RUI2CTpop workflows, we recommend computational hardware with 10 or more cores, 256GB RAM, and at least 4TB of disk space. We recommend using a Linux-based environment like Ubuntu as the operating system of choice. For the DCTA workflow, annotations per dataset can be distributed among similarly configured servers. In our cluster, we used five Linux servers running 10 dataset annotation runs per server in parallel for a maximum of 50 runs executing in parallel.

## Data used

A table on GitHub^99^ lists dataset IDs for all 16,293 datasets originally downloaded.

### Data portals

#### HuBMAP

As of December 2025, the HuBMAP Data Portal (portal.hubmapconsortium.org) listed 4,902 public datasets from 27 organs. These datasets were ingested by tissue providers through a UI (ingest.hubmapconsortium.org) where they could enter donors, organs, samples/tissue blocks, tissue sections, etc. Relationships between these entities are organized in a provenance hierarchy where a donor and organ are needed so that tissue samples can be organized based on diverse tissue sample types. APIs enable users to access entities programmatically (docs.hubmapconsortium.org/apis.html). Both published and unpublished datasets exist on the HuBMAP and SenNet portals (see below). Published datasets have been sent through a series of QA/QC processes. Unpublished datasets are only accessible with authentication. Only published datasets were used for HRApop construction. Concatenations of HuBMAP data organized by assay and organ exist at data-products.hubmapconsortium.org/data_products.

#### SenNet

The SenNet Data Portal (data.sennetconsortium.org/search) uses a similar infrastructure as the HuBMAP Data Portal. It features human and murine datasets. As of December 2025, 1,174 human datasets from 214 donors were publicly available. Like for HuBMAP, APIs provide programmatic access to datasets, donors, organs, etc., see docs.sennetconsortium.org/apis.

#### GTEx

The GTEx Portal (www.gtexportal.org/home/downloads/adult-gtex) hosts the adult GTEx data and resources and provides open access to, e.g., expression quantitative trait loci (eQTLs), and protected access, e.g., to limited donor phenotypes as well as de-identified donor data for sequencing. For GTEx sc-data, one H5AD file with data for all 8 organs plus donor metadata was downloaded (storage.googleapis.com/adult-gtex/single-cell/v9/snrna-seq-data/GTEx_8_tissues_snRNAseq_atlas_071421.public_obs.h5ad).

#### CxG

The CxG Portal (cellxgene.cziscience.com/collections) provides access to both primary and secondary datasets. Primary datasets contain the raw or minimally processed data while secondary datasets are curated and normalized. The DCTA Workflow retrieved the secondary datasets when possible unless they contained fewer data points than the primary datasets. As of December 2025, there were 1,015 collections from healthy donors older than 15 years.

#### Cell counts

Cell counts represent the unprocessed number of RNA transcripts detected for each gene in each cell. To ensure data integrity and consistency, raw cell counts were used wherever available during CTann. For HuBMAP and SenNet datasets, these were obtained from the “counts” layer of the H5AD file, which contains integer counts per gene and cell, while for GTEx and CxG datasets, raw counts were obtained from the “raw.X” attribute of the H5AD file, which stores original and unnormalized counts for each cell and gene.

#### Crosswalks

To enable comparisons between CTs assigned by different CTann tools (and sc-proteomics data, which used human-assigned CTs), CT labels needed to be crosswalked to CTs in the anatomical structures, cell types, plus biomarkers (ASCT+B) tables^36^ using CL or PCL terms. Crosswalks for each CTann tool were curated manually by experts and are published at lod.humanatlas.io/ctann; the underlying HRA Digital Object type is described in a related publication^22^. Since not all CTs had an exact match to a CL term, the Simple Knowledge Organization System (SKOS, www.w3.org/2004/02/skos)^100,101^ was used to indicate if the mapping was done to a term with an exact match (skos:exactMatch) or if they had to be mapped to a more general class (skos:narrowMatch). The crosswalks published in HRA v2.3 linked 1,615 annotation labels and 1,909 annotation IDs from Azimuth, CellTypist, popV, and sc-proteomics datasets (author-assigned) to 885 CL labels and 495 CL IDs for 36 organs. 1,923 mappings were exact matches and 683 were narrow matches.

The DCTA Workflow applied crosswalks after CT annotations were done; CL labels and IDs were used when computing CT populations for ASs, datasets, and extraction sites. Exactly 154 unique cell labels could not be crosswalked to CL and a report of modality, CTann tool (if applicable), organ, and CT label for these in **Table S1**.

## Existing code

### CTann tools

The three CTann tools in HRApop v1.0 (Azimuth^49^, CellTypist^50,51^, popV^52^) were containerized with Docker in the HRApop CTann Tool Containers on GitHub^94^. The Dockerfile listed operating system requirements, basic setup, and dependencies so the package could be run in the cloud or on a local machine with consistent outputs. This enabled the DCTA Workflow to execute them locally. These containers defined a dataset handler interface (see code on GitHub^102^) that specified requirements for every new piece of code that downloaded H5AD files from a portal. Docker was used to package each CTann tool along with all of its dependencies into portable containers that could be deployed across different environments. Apptainer (apptainer.org), was used when running HRApop code on HPC clusters. Each container had a context file that specified which models needed to be downloaded, such as this one^103^ for Azimuth. Similarly, extracting summaries, computing gene expressions, and crosswalking were also packaged as Docker files. Both Docker and Apptainer/Singularity support running Common Workflow Language (CWL, www.commonwl.org) workflows on a Linux cluster for HRApop v1.0. An example for Azimuth is available on GitHub^104^. **Table S3** lists the name, version number, code base, models used, and requirements for each tool.

### Mean biomarker expressions for sc-transcriptomics data

Mean biomarker expressions were captured in the DESpop and provided for each CT per dataset. To generate these values, *scanpy*^105^, *numpy*^106^, and *anndata*^107^ were used. Specifically, *scanpy*’s *rank_gene_groups()* method (scanpy.readthedocs.io/en/stable/generated/scanpy.tl.rank_genes_groups.html) was applied to perform differential expression analysis between CTs. As part of this analysis, this method calculated the mean expression of each gene within a target CT, as well as its expression in the rest of the dataset. These values were used to identify and rank marker genes, and the corresponding mean expressions were recorded for the top-n genes, where n was defined by the user. However, it is important to note that this method does not compute mean expression values for all genes across all CTs—only for the most differentially expressed ones. To ensure consistency in gene naming across datasets, gene identifiers were normalized using a lookup table, which maps Ensembl IDs from Release 111^108^ (www.ensembl.org/index.html) to HGNC-approved symbols from version v2023-09-18^109^ (www.genenames.org). This normalization helps maintain interoperability and accuracy when comparing gene expression data across datasets and tools.

### Tissue registration

Registering tissue datasets inside the 73 3D reference objects in the HRA was made possible via the RUI^24^, which generates cuboid, 3D extraction sites; it is available as a standalone tool at apps.humanatlas.io/rui but also embedded into the ingest pipelines of HuBMAP and SenNet. The registration process consisted of three main phases: assignment, enrichment, and validation. Initially, the registration coordinator contacted subject matter experts with knowledge of the spatial information associated with tissue samples. Experts could then use the RUI to submit spatial information for tissue blocks themselves or collaborate with the coordinator, who facilitated the submission process with their input. The RUI recorded spatial information by creating extraction sites inside 3D reference objects. Additionally, it used mesh-based collision detection to annotate the extraction with AS tags (see **Box 1** and **Mesh-based collision detection** section). By the end of this phase, all tissue samples had an assigned extraction site. These workflows are further detailed in SOPs^26–28^.

Once extraction sites were assigned, enrichment began. The registration coordinator used a location processor tool to enhance spatial information with de-identified donor metadata (e.g., sex, age, BMI) and publication metadata (e.g., DOI, authors, publication year). This combined dataset formed a registration set^35^, which was assigned a unique ID for future reference. The code for the processor is accessible via GitHub^110^, as are existing stand-alone HRA Registrations^91^.

Finally, in the validation phase, the expert was asked to review the registration set for accuracy and completeness, requesting revisions if necessary. This process uses a customized instance of the EUI^24^, which the expert used to evaluate sample locations, metadata accuracy, and AS tags from mesh-based collision detection. Once validated, the registration set was finalized and added to the general EUI (apps.humanatlas.io/eui). This concluded the spatial registration process. An overview of all EUIs for registration sets is available on GitHub^111^.

### Mesh-based collision detection

To enable more precise collision detection between tissue blocks and ASs, a library for mesh-based collision detection (see **Box 1**) was created and named HRA Mesh Collision API^112^. Given an extraction site, this HTTP service returns a list of mesh collisions with ASs and metadata. The 3D geometry-based tissue block annotation code includes: (1) a C++ library for the HTTP service for collision detection and intersection volume (see **Box 1**) computation between extraction sites and ASs, (2) a C++ library for checking manifoldness and closedness of meshes as well as hole-filling for unclosed meshes, and (3) a Python library for converting GLB files to Object File Format (OFF) files, used as the underlying 3D model format for collision detection. The code repository, URL to deployed API, and exemplary API response are available in **Table S1**.

### Weighted cosine similarity

A collection of functions to compute and use weighted cosine similarities in the RUI2CTpop Workflow is available on GitHub^113^. The script uses math.js (mathjs.org) for access to implementation for the dot product and norm between two vectors (mathjs.org/docs/reference/functions/dot.html and mathjs.org/docs/reference/functions/norm.html).

## New code

### HRApop CTann Tool Containers

These were a collection of scripts (see GitHub^94^) for annotating H5AD files using Azimuth^49^, CellTypist^50,51^, and popV^52^. They wrapped CTann tools as Docker containers for CWL workflows. This allowed each tool to be executed consistently regardless of technology needed, e.g., R vs Python. Default settings were used for each tool where possible. A complete list of settings for each tool is available in **Table S3.**

### DCTA Workflow

Downloading and annotating datasets was handled by the **DCTA Workflow** (see GitHub^93^). First, it ensured that the data from multiple portals was from healthy, human donors, then ran applicable CTann tools via the **HRApop CTann Tool Containers**, analyzed gene expressions to identify top genes with *scanpy*, crosswalked CT labels from CTann to ASCT+B tables^36^ using crosswalks, assembled donor metadata, and output summarized results for downstream use. It was runnable as a CWL workflow. The CWL runner was written in Python. The DCTA Workflow then produced CT populations and metadata for all annotated sc-transcriptomics and sc-proteomics datasets as output (see **Table S1**). They were then copied to the input GitHub repository^114^ for the RUI2CTpop Workflow (see below) for further processing.

### Download

To download H5AD files, the DCTA Workflow constructed a series of jobs to execute. An organ mapping provided crosswalks between organ code names on the portals to Uberon IDs and labels on HuBMAP and SenNet (see this GitHub commit^115^) as well as GTEx (see this GitHub commit^116^). For CxG, no mapping was needed as the metadata contained Uberon IDs already. To retrieve donor metadata across portals, each portal was queried according to their APIs, then relevant information was extracted and saved in a harmonized format, see donor field at this GitHub commit^117^. Implementations to extract donor metadata from the different portals are also available, e.g., for SenNet (see this GitHub commit^118^) and CxG (see GitHub^119^).

The DCTA Workflow extracted metadata needed for constructing ds-graphs^22^ (age, sex, BMI, assay type) from the individual portal APIs and saved it as JSON files. The H5AD files were downloaded locally into a raw data folder in the DCTA Workflow repository, or onto a file system on a HPC system.

### Splitting and re-assembling H5AD files for GTEx and CxG data

In the data model of HuBMAP and SenNet, donors, organs, tissue blocks, tissue sections, and datasets are modeled as individual entities, where each dataset belongs to exactly one donor. This means that an H5AD file from HuBMAP or SenNet contains data for exactly one donor. On CxG, on the other hand, H5AD files contain multiple donors; to make the two data models work together, H5AD files from CxG needed to be split into new H5AD files by donor and organ in the DCTA Workflow. The respective script^120^ was written in Python, because it needed *pandas*, a foundational library for data manipulation and analysis (pandas.pydata.org), and *anndata* for handling annotated data matrices (anndata.readthedocs.io/en/stable). This was done to combine donor-organ combinations across assets into new H5AD files. Extracted donor metadata fields were shown in the harmonized donor metadata format described above. These new H5AD files are made available alongside HuBMAP and SenNet H5AD files on Globus^121^.

### Datasets with too few cells

Datasets with fewer than 100 cells were filtered out by the DCTA Workflow. While their H5AD files were downloaded, no CTann was run over them and no CT population was output.

#### RUI2CTpop Workflow

The **RUI2CTpop Workflow** (see GitHub^96^) performed spatial annotation and CT population computation with input files provided by the DCTA Workflow. It sourced extraction sites via the HuBMAP and SenNet APIs through HRA API queries at apps.humanatlas.io/api#get-/ds-graph/hubmap, apps.humanatlas.io/api#get-/ds-graph/sennet, and apps.humanatlas.io/api#get-/ds-graph/gtex, with the underlying queries at github.com/x-atlas-consortia/hra-api/blob/main/src/library/ds-graph/operations/hubmap.js, github.com/x-atlas-consortia/hra-api/blob/main/src/library/ds-graph/operations/sennet.js, and github.com/x-atlas-consortia/hra-api/blob/main/src/library/ds-graph/operations/gtex.js. A listing of all extraction sites that are sources is available at github.com/x-atlas-consortia/hra-pop/blob/main/input-data/v1.0/config.sh.

For the RUI2CTpop Workflow to function, the DCTA Workflow provided CT populations and dataset metadata, then those files were copied over to the input folder for a new RUI2CTpop run (see GitHub^69^). Scripts running over these input files during RUI2CTpop are on GitHub^122^. Output data from **RUI2CTpop** is provided on GitHub^123^.

THe RUI2CTpop Workflow processed Criteria C1-4 (including a check for donor age to ensure only data from adult humans is used, see **Box 1**) and gathered extraction sites, CT populations, donor metadata, and related publications where applicable. If a dataset had no metadata for age or sex, it was not used for atlas construction. GTEx provided only age ranges, not values, but the data came from adult donors. The RUI2CTpop Workflow then used mesh collisions to build the ASpop and the DESpop (see both definitions in **Box 1**, links in **Table S1**) and corridor code to compute corridors (see **Corridors** section). Exemplary CT populations for a dataset, an extraction site, and an AS are shown at cns-iu.github.io/hra-cell-type-populations-supporting-information/#exemplary-cell-type-populations. Note that in all three cases, the CTann tool(s) are indicated by the annotation_method field. The RUI2CTpop Workflows also contained scripts and SPARQL queries to construct data products for HRApop in the form of CSV reports^92^ to analyze, visualize, validate, and use HRApop data (see link in **Table S1**).

### Corridors

For each extraction site with a CT population, a 3D volume of likely origin within the organ was computed, given the biomolecular make-up of the tissue block as represented by its CT population. The result was a complex corridor, i.e., a combined representation for all possible locations, compiled via a shrink-wrap algorithm^124^. Corridors represented the complete set of spatial positions where extraction sites could plausibly be located while maintaining their observed intersection ratios with neighboring ASs. Each extraction site was uniquely associated with one such corridor. Corridors were GLB files (see **Data Records**). The spatial origin could be an entire AS if it had the same or a similar CT population (measured using weighted cosine similarity) or the extraction site of a tissue block with the most similar CT population (and its corresponding corridor with the same percentages of multiple ASs).

To generate complex 3D corridors given an extraction site with the RUI, a C++ library with an HTTP service for the 3D Corridor Generation API^125^ (apps.humanatlas.io/api/#post-/v1/corridor, see also **Table S1**) was created. Corridors were computed by sending an extraction site to this API. From there, three cases are possible: (1) If the extraction site collided with **only one AS**, the entire AS was returned as a corridor; (2) if the extraction site collided with **two ASs,** a filter-search algorithm was used to efficiently compute all the possible locations before applying a shrink-wrap algorithm to generate a complex corridor. The filter-refine paradigm^126^ is widely used in computationally intensive tasks where infeasible solutions are filtered out from a list of candidates. Next, more viable candidates are examined with respect to their exact geometry to generate exact answers in a refinement step. Inspired by the filter-refine paradigm, a filter-search algorithm was made to derive complex corridors. Finally, (3) if a tissue block collided with **three or more ASs,** it was fixed in place, in which case it corresponded exactly to the extraction site.

Corridors were made available as a ZIP file on Zenodo^82^; they were named after the extraction site on which were based, e.g., corridors/1cbd9283-2d58-4a2d-88fe-effb18c3f14f.glb, which belongs to the extraction site with ID 1cbd9283-2d58-4a2d-88fe-effb18c3f14f from the head of the female pancreas. I can be inspected in 3D on the companion website at cns-iu.github.io/hra-cell-type-populations-supporting-information#exemplary-corridor. Corridors were captured in the binary Graphics Library Transmission Binary Format (GLB, www.khronos.org/gltf). HRApop made 1,189 corridors available, including the 230 corridors for the 230 extraction sites in DESpop, plus 959 corridors for extraction sites not in the HRApop Atlas. Their total size was 202 MB (99.8 MB when compressed). Code, endpoint, and documentation for the 3D Corridor Generation API is available in **Table S1**.

### Tradeoff between step size in search stage and precision of corridor

Since a sliding window approach was used to search feasible locations if two or more AS collided with an extraction site, the configuration of the step size was essential. It determined how big a move was made in the search stage (see **Table 2**). If a large step size was set, locations with the exact intersection volume (see **Box 1**) with the given extraction site could be skipped. Conversely, if a small step size was set, the computation cost could become overwhelming. Further, the intersection volume from the mesh-based collision detection was returned as a float; thus, if only the exact value was matched, there could be very few or no feasible locations. In order to compute corridors with both high precision and reasonable computation overhead, the tolerance for feasible locations had to be adjusted to, e.g., 0.1, which means the difference between the intersection volume of the feasible locations and the true intersection volume cannot exceed 10% of the true intersection volume.

**Table 2.** Filter and search stage for pre-computing corridors.

|  |  |  |
| --- | --- | --- |
| 1 | **Filter stage** | Each collided mesh was approximated by their minimum bounding boxes. Then, the area $\Omega$ where a fixed-size, axis-aligned tissue block could be put was computed so that it could intersect with all the minimum bounding boxes of meshes. |
| 2 | **Search stage** | A brute force search algorithm was applied by specifying the step size. A sliding window approach was used to move the fixed-size tissue block within the searching area $\Omega$ computed in the filter stage. |

## Data Records

The dataset is available on Zenodo (zenodo.org, processed data and archived code), with raw data on Globus (www.globus.org) and actively developed code on GitHub.

### HRApop data products on Zenodo

Six major HRApop data products are available for download on Zenodo^82^:

1. Atlas (covers 73 ASs in 17 organs via 662 HRApop Atlas datasets: 558 sc-transcriptomics, 104 sc-proteomics)
  a. ASpop (as JavaScript Object Notation for Linked Data format, or JSON-LD, see json-ld.org).
  b. DESpop (JSON-LD).
2. Input for the RUI2CTpop Workflow.
  a. Cell instances with top 10k genes for all datasets downloaded for the DCTA Workflow (compressed CSV).
  b. Corridors for all extraction sites used as input for the DCTA Workflow (ZIP folder with GLB files). Contains 1,189 corridors, including 230 corridors in DESpop.
3. QC
  a. Cell instances (see **Box 1**) with confidence scores for each CTann tool assignments (compressed CSV), used for **Fig. 4**.
  b. QC metrics for all sc-transcriptomics datasets that are annotated via the DCTA Workflow are provided as a ZIP file with one folder per dataset, used for **Table 5** in the **Technical Validation**.

### Direct download links for CSV files

**Table 3** below provides links to download ASpop and DEspop as CSV files.

**Table 3.** Direct links to CSV files for ASpop and DESpop.

| Name | URL | Type |
| --- | --- | --- |
| CT populations for ASs | apps.humanatlas.io/api/grlc/hra-pop/cell_types_in_anatomical_structures_per_as.csv | ASpop |
| CT populations for extraction sites | apps.humanatlas.io/api/grlc/hra-pop/cell-types-per-extraction-site.csv | DESpop |
| CT populations for datasets | apps.humanatlas.io/api/grlc/hra-pop/cell-types-per-dataset.csv | DESpop |

### Raw data on Globus

All H5AD files used for construction ASpop and DESpop for sc-transcriptomics and sc-proteomics are provided for download on Globus^121^.

### Archived code

The release code for HRApop v1.0 was archived on Zenodo for the DCTA Workflow^127^, the HRApop CTann Tool Containers^128^, and the RUI2CTpop Workflow^129^.

### Miscellaneous

A full listing of repositories used to construct and use HRApop v1.0 is provided in **Table S1**. Examples for usage of this HRApop Atlas Data are available on the companion website at cns-iu.github.io/hra-cell-type-populations-supporting-information#usage-examples. The **Usage Notes** section details how to access HRApop data via Jupyter Notebooks.

## Data Overview

### Counts for HRApop v1.0

On June 16, 2025, the RUI2CTpop Workflow was run to compile HRApop v1.0. It downloaded 16,293 datasets with 57,911,931 cells from the four sc-portals, which were then sent through a filtering process. Additionally, as of December 2025, 1,746 tissue blocks from 1,327 different 3D cuboid extraction sites exist across 30 organs and they link to 6,378 tissue datasets. 558 of the 662 datasets in the HRApop Atlas Data are sc-transcriptomics datasets that were annotated using Azimuth^49^, CellTypist^50,51^, and/or popV^52^, covering 11,042,750 unique cells in a total 3D volume of ∼12.05 liters (dm³) with partially intersecting extraction sites in 73 unique ASs in 17 organs (112 ASs and 31 organs if male and female are counted separately). The datasets come from 230 3D extraction sites that cover 54 3D ASs across 17 organs. While the HRApop Atlas focuses on these 558 sc-transcriptomics datasets in 17 organs, the method is generalizable to sc-proteomics (see **Generalization to spatial data** section) and to all organs. **Table 4** provides counts of datasets and cells in the HRApop Atlas Data, split by sex, consortium, number of datasets, number of cells, and modality. HuBMAP, SenNet, and GTEx have their own portals. HCA and NHLBI/LungMap^130,131^ datasets all come from the CxG Portal. Detailed counts for the HRApop Atlas from the RUI2CTpop Workflow are provided in **Table S4.**

**Table 4.** HRApop Atlas Datasets. A breakdown of datasets that meet Criteria C1-4 (see **Box 1**) and are used to construct HRApop v1.0.

| Sex | Consortium | #Datasets | #Cells | Modality |
| --- | --- | --- | --- | --- |
| Female | GTEx | 7 | 47,863 | sc_transcriptomics |
| Male | GTEx | 8 | 70,113 | sc_transcriptomics |
| Female | HCA | 63 | 364,993 | sc_transcriptomics |
| Male | HCA | 58 | 359,273 | sc_transcriptomics |
| Female | HuBMAP | 22 | 900,547 | sc_proteomics |
| Female | HuBMAP | 112 | 2,933,015 | sc_transcriptomics |
| Male | HuBMAP | 82 | 15,676,316 | sc_proteomics |
| Male | HuBMAP | 253 | 6,626,620 | sc_transcriptomics |
| Female | NHLBI/LungMap | 1 | 4,680 | sc_transcriptomics |
| Male | NHLBI/LungMap | 2 | 5,713 | sc_transcriptomics |
| Female | SenNet | 18 | 225,940 | sc_transcriptomics |
| Male | SenNet | 36 | 404,540 | sc_transcriptomics |
| **TOTAL** |  | **662** | **27,619,613**<br>sc-transcriptomics: **11,042,750**<br>sc-proteomics: **16,576,863** |  |

### Anatomical Structure Cell Type Populations

**Fig. 3** shows the 112 ASs of the male (left) and female (right) reference body for which spatially registered sc-transcriptomics data existed in HRApop v1.0. For each AS, the organ name, number of datasets, and AS name plus a bar graph with the percentage of major CTs, i.e., ASpop are shown (see **Box 1**). Only sc-transcriptomics data is shown and Azimuth^49^, CellTypist^50,51^, and/or popV^52^ used to annotate cells; with a preference for Azimuth annotation over CellTypist over popV^52^. Crosswalking to CL or PCL and aggregation to higher-level CTs is also described in detail in the **Crosswalks** section. For the left and right mammary gland, two CTs that cannot be mapped to a high-level CT make up ∼70% of the CTs across ASs. As a result, the stacked bar graphs for these are mostly grey: luminal epithelial cell of mammary gland (purl.obolibrary.org/obo/CL_0002326) with ∼53% and fibroblast of breast (purl.obolibrary.org/obo/CL_4006000) with ∼16%.

**Fig. 3.**
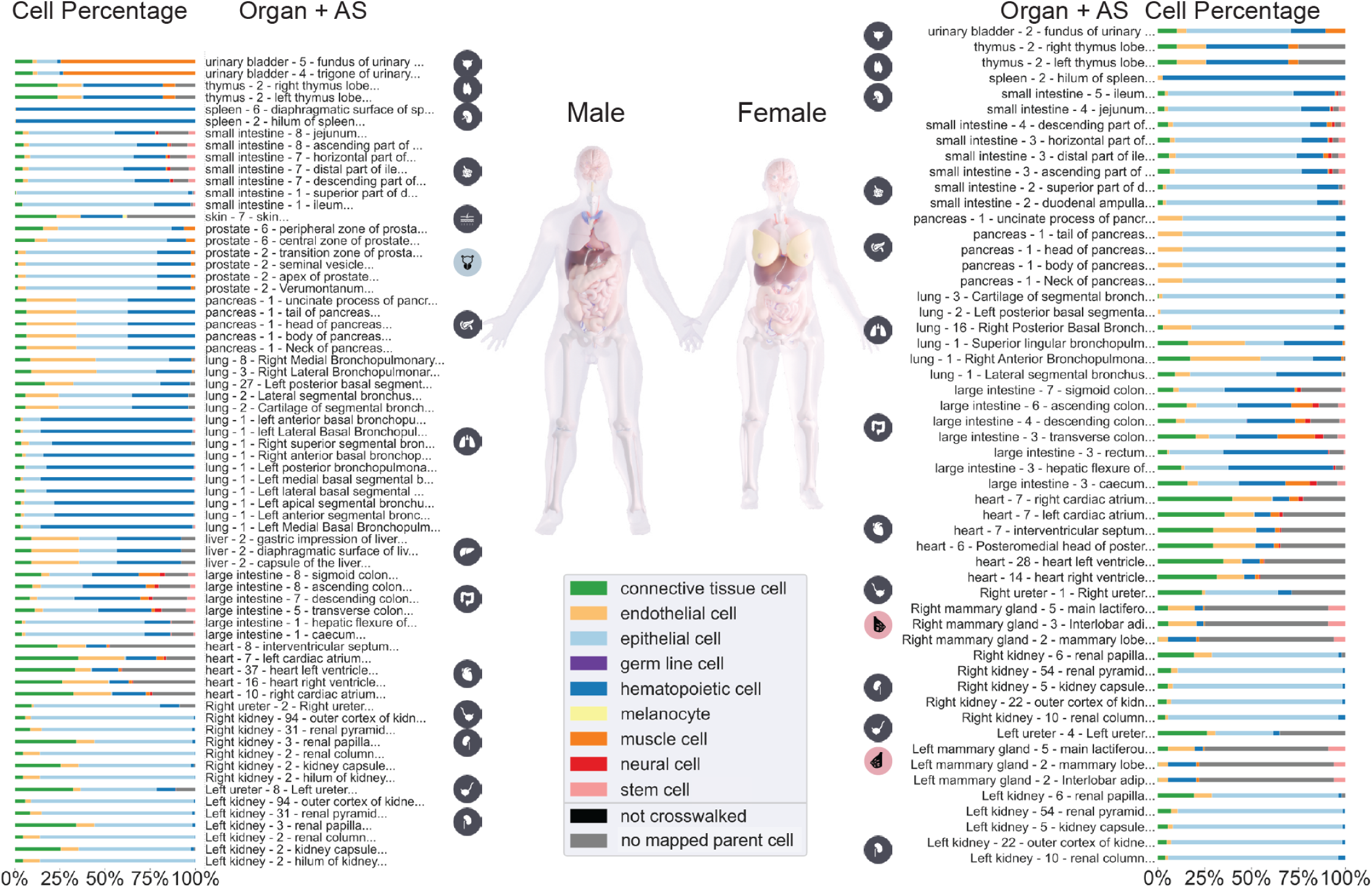
CT populations for unique ASs across male and female in HRApop v1.0. Stacked bar graphs of the percentage of CT identified in ASs, aggregated to higher level CTs from CL, see listing on GitHub^132^, shown for male (left) and female (right, placenta was omitted). The male-only organ icon (prostate) was rendered in blue while icons for female-only organs were rendered in pink (mammary gland left and right).

Data featured in **Fig. 3** used CTann crosswalks to harmonize and compare datasets across portals. The resulting CT typology has 201 low-level CTs. CL defines 11 broad, high-level classes — a set curated by CL editors to cover most low-level CTs while minimizing overlap; it is available on GitHub^133^. The 201 cell types in HRApop fall into only 9 of these classes, leaving two unpopulated (extraembryonic cell and bone cell). For visualization purposes, the 201 types were grouped into these 9 populated high-level classes. Additional mappings exist to 19 medium-level, more granular CTs—a subset of terms that, while still broad, provides a more detailed classification—via CL IDs and labels; this mapping is available in **Table S1**. The mapping applied in **Fig. 3** used the 9 CTs belonging to the upper slim of high-level CTs directly from CL (see GitHub^133^); note that CTs from PCL are included in this slim ontology 26 CTs were classified under more than one category in CL; for these, one of the classes was chosen reflecting expected biological grouping (see **Table S5**).

If a CT was marked as “no mapped parent cell,” the CT term (already crosswalked to CL or PCL) was not a subclass of a top-level CT in the top-n CTs provided by CL; in HRApop v1.0, 617,000 cells associated with 10 CTs are rendered in gray. If a CT was marked as “not crosswalked,” this means that the CT label assigned by the CTann tool was not associated with a CL or PCL ID.; in HRApop v1.0, 9 cells associated with 1 CT in the small and large intestine that were not covered in existing ontologies and are rendered in black. A full report of cells that were not crosswalked while constructing HRApop but that have been crosswalked semi-manually for **Fig. 3** was labeled “unmapped-cell-ids” and is available on GitHub^134^.

## Technical Validation

Four validations are presented in this section: for sc-transcriptomics data, we show (1) bins of confidence scores for each cell instance by tool, mean, and median ribosomal plus (2) mitochondrial gene percentages with standard deviation; for sc-transcriptomics and sc-proteomics, we show (3) heatmaps with different CT prevalence between AS in the same organ and (4) the number of datasets per AS by organ.

### Sc-transcriptomics data

#### Confidence scores per cell per tool

When running the DCTA Workflow, Azimuth, CellTypist, and popV computed confidence scores for each cell instance annotation. **Fig. 4** shows a histogram where the x-axis shows confidence scores (with bin width 0.002), the y-axis shows the number of cells per bin, and color encodes the tool assigning the confidence scores. The mean confidence scores are: Azimuth (mean=0.62, median=0.68, SD=0.26), CellTypist (mean=0.46, median=0.34, SD=0.37), popV (mean=0.71, median=0.67, SD=0.24). The histogram shows that Azimuth and CellTypist have different values for their measures of central tendency while both generating continuous confidence scores, whereas popV generated spikes due to its averaged voting mechanism For this paper, each tool has their own strengths and weaknesses, and as related work on benchmarking different CTann tools has shown^76^, when there is a discrepancy, there is no consensus on the best tool. This Data Descriptor provides results of a scalable, reproducible workflow that runs three CTann tools at scale while providing the user with data products documenting the results so that they can apply their expertise to assess the CTann assignments. The code to reproduce the histogram in **Fig. 4** is provided on GitHub^135^. The compressed CSV file is available on Zenodo^82^ (filename: “sc-transcriptomics-cell-instances.csv.gz”).

**Fig. 4.**
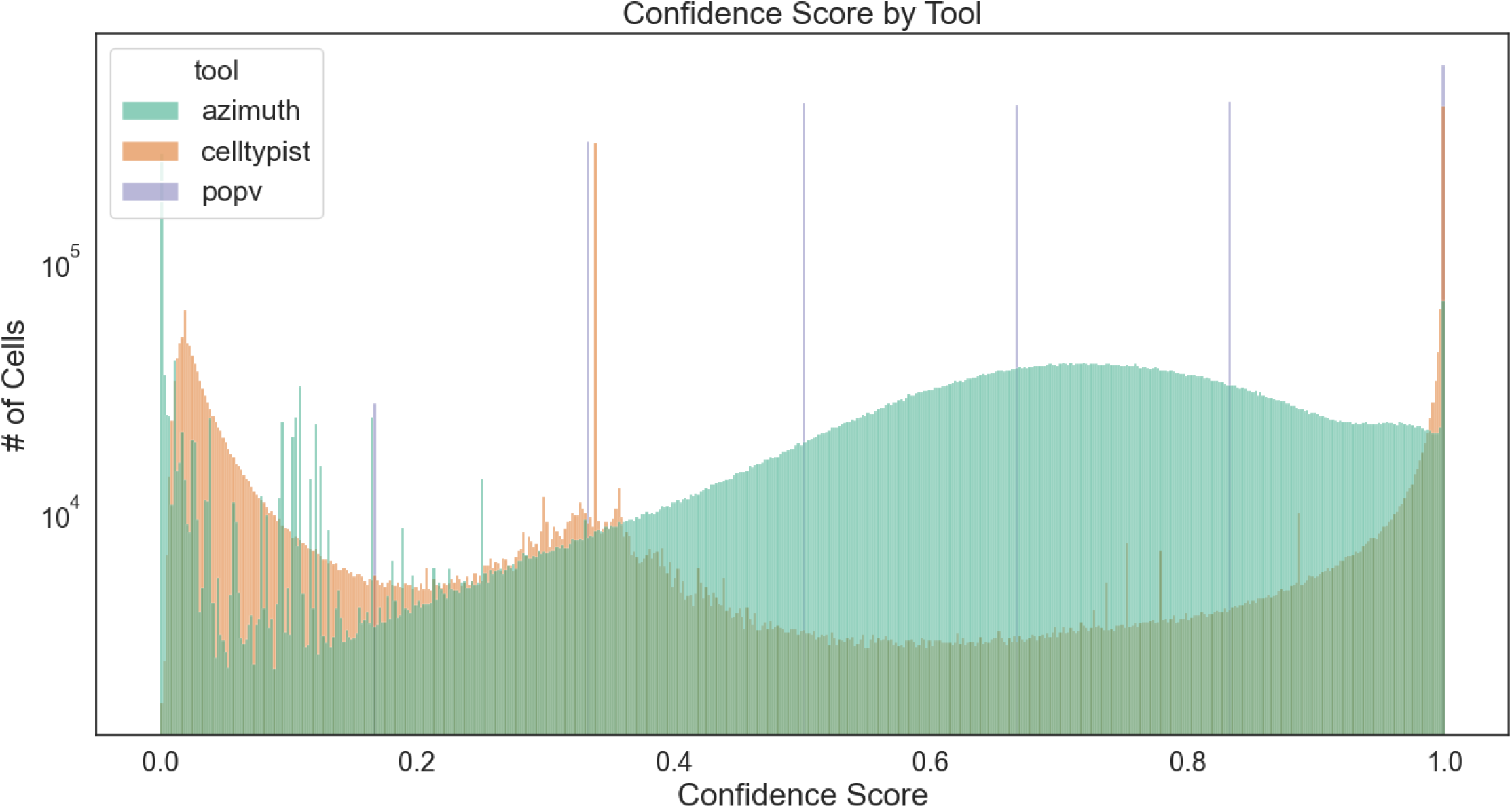
Histogram of confidence scores by number of cells with that score, using a bin width of 0.002, colored by CTann tool.

#### Ribosomal and mitochondrial gene percentages per organ

Using *scanpy*’*s* built-in *calculate_qc_metrics()* function (scanpy.readthedocs.io/en/stable/generated/scanpy.pp.calculate_qc_metrics.html), mean and median ribosomal and mitochondrial gene percentages for all 558 sc-transcriptomics datasets (H5AD files) were computed and then aggregated to the mean and median at the organ level plus standard deviation, see **Table 5**. For QC purposes, the threshold for mitochondrial gene percentages for single-nucleus and single-cell data is different (less than 2% for single nucleus, less than 10% for single-cell). Thresholds for ribosomal genes are less standardized than mitochondrial percentage, but there are practical norms used in QC (5–20% ribosomal for single-nucleus, 20–40% ribosomal content is common for single-cell). While the available metadata from HuBMAP and SenNet made it impossible to reliably capture whether a dataset was single-nucleus or single-cell, **Table 5** shows that all mean and median percentage values are well below any of the single nucleus and single-cell thresholds. Mitochondrial percentages are 0 across all organs and metrics. For ribosomal percentages, the largest individual value is the median percentage in the male reproductive system at 0.0015. This is to be expected, given that only datasets from portals with built-in QA/QC were used or datasets with an associated peer-reviewed publication.

**Table 5.** Mean and median ribosomal and mitochondrial gene percentages across datasets per organ with standard deviations.

| Organ name | Mean percentage ribosomal genes | Median percentage ribosomal genes | SD percentage ribosomal genes | Mean percentage mitochondrial genes | Median percentage mitochondrial genes | SD percentage mitochondrial genes |
| --- | --- | --- | --- | --- | --- | --- |
| Set of lactiferous glands in left breast | 0 | 0 | 0 | 0 | 0 | 0 |
| Set of lactiferous glands in right breast | 0.00133 | 0.001 | 0.00058 | 0 | 0 | 0 |
| Heart | 0.00005 | 0 | 0.00028 | 0 | 0 | 0 |
| Large intestine | 0 | 0 | 0 | 0 | 0 | 0 |
| Left kidney | 0 | 0 | 0 | 0 | 0 | 0 |
| Left ureter | 0 | 0 | 0 | 0 | 0 | 0 |
| Liver | 0 | 0 | 0 | 0 | 0 | 0 |
| Male reproductive system | 0.00117 | 0.0015 | 0.00098 | 0 | 0 | 0 |
| Pancreas | 0 | 0 | 0 | 0 | 0 | 0 |
| Respiratory system | 0.00009 | 0 | 0.00056 | 0 | 0 | 0 |
| Right kidney | 0 | 0 | 0 | 0 | 0 | 0 |
| Right ureter | 0 | 0 | 0 | 0 | 0 | 0 |
| Skin of body | 0.00071 | 0 | 0.00144 | 0 | 0 | 0 |
| Small intestine | 0 | 0 | 0 | 0 | 0 | 0 |
| Spleen | 0 | 0 | 0 | 0 | 0 | 0 |
| Thymus | 0 | 0 | 0 | 0 | 0 | 0 |
| Urinary bladder | 0 | 0 | 0 | 0 | 0 | 0 |

A ZIP file with the QC results containing one directory per dataset. We provide the raw QC results that this table and figure are available on Zenodo^82^ (filename: “hra-pop-v1.0-qc.zip”). The code to reproduce **Table 5** is provided on GitHub^136^.

### Sc-transcriptomics and sc-proteomics data

#### Heatmaps

While some CTs exist across organs (e.g., macrophages), most CTs are highly specialized to deliver well-defined organ-specific functions in ASs. To demonstrate that ASpop varies not only by organ but also by AS, four heatmaps were made (one per CTann tools plus sc-proteomics, see **Fig. C1** and **Fig. C2**; high-resolution versions are available on the companion website at cns-iu.github.io/hra-cell-type-populations-supporting-information#figures. Each heatmap lists CT labels on the x-axis and organ plus AS labels on the y-axis. Table field color represents the scaled mean value, i.e., z-score, see **equation (1)** for the percentage of CTs identified in each AS.

For each heatmap, data from a CTann tool is selected and processed to calculate the average CT percentage associated with all the ASs in an organ. The results are transformed from a data frame into a matrix (CTs by organ plus AS concatenated into a combined label), where each matrix cell represents the average CT percentage measured for CT and AS dyad. Finally, matrix values are converted to a standardized z-score, which is calculated using the formula in **equation (1)**.

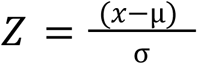

**Equation 1. Computation of standardized z-score for CTs across ASs.**

**Equation (1)** shows *x* as an average CT percentage measure, *μ* is the mean average CT percentage, and *σ* is the standard deviation mean average CT percentage. The z-score identifies how many standard deviations a data point is from the average mean. If the z-score is 0, values are close to the variable’s average; a z-score of 1 indicates that CT percentage values are 1 standard deviation higher than the mean for that CT, values of 2 are 2 standard deviations from the mean, etc.

Differences across organs and CTann become visible. For example, heart, lung, kidney, and pancreas, when annotated with Azimuth (see **Fig. C1A**), show distinct bands of CTs with a z-score of ∼1.5, some up to 5. We observe similar patterns for the heart, liver, lung, pancreas, skin, and small and large intestines in CellTypist (see **Fig. C1B**) and breast, heart, liver, lung, pancreas, prostate, skin, small and large intestines, spleen, thymus, urinary bladder, and ureter for popV (see **Fig. C1.C**). High-resolution versions of the heatmaps and the code to generate them are listed on the companion website at cns-iu.github.io/hra-cell-type-populations-supporting-information#z-scores-for-cts-per-organ-and-as.

#### Datasets per AS

The grouped bar graphs in **Fig. 5** show the coverage of HRApop Atlas Data across ASs and organs. Depicted is the number of datasets on the x-axis per AS on the y-axis (labeled by organ for better groupings). Overlaps and gaps in coverage between CTann tools (horizontal), across all organs and ASs (vertical) and between sex (color) become visible. The AS with the most registered HRApop Atlas Data is the outer cortex of the kidney (purl.obolibrary.org/obo/UBERON_0002189), with 75 datasets run through Azimuth. Sc-proteomics datasets are omitted. Counts for this graph are provided on the companion website at cns-iu.github.io/hra-cell-type-populations-supporting-information#dataset-counts.

**Fig. 5.**
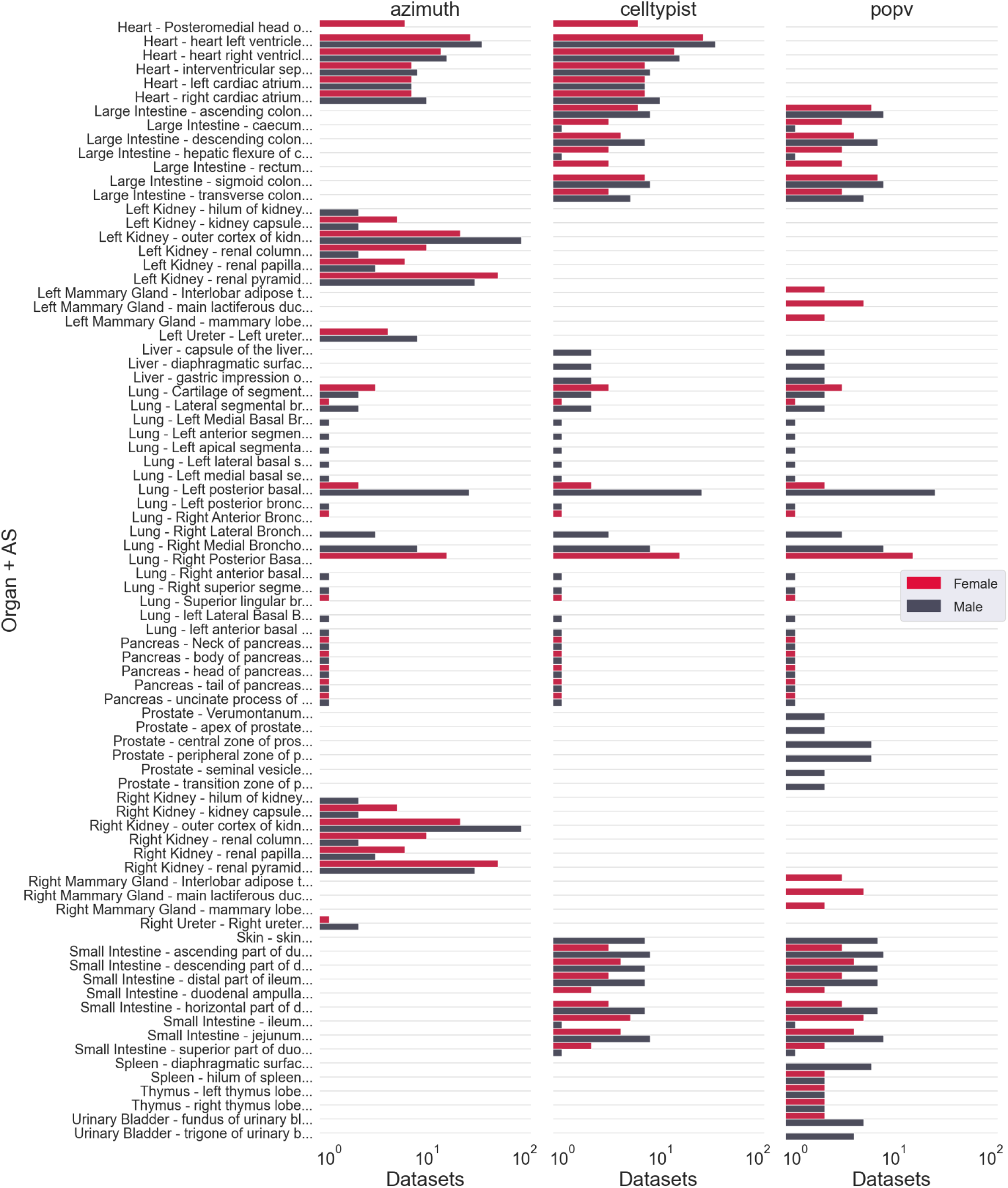
HRApop Atlas Datasets per sex-specific organ and AS. The number of HRApop Atlas Data per organ and AS combination (y-axis) is plotted on the x-axis, faceted by tool (columns) and colored by sex. Note that the x-axis is scaled logarithmically.

## Usage Notes

The **Data Records** section points to download links for 558 H5AD files for sc-transcriptomics data, CT populations with expert-provided CTann for sc-proteomics data, and HRAop data products. This section details two additional ways of accessing processed HRApop data.

### Getting HRApop data via API queries

A Jupyter Notebook detailing programmatic access to CT populations for ASs, extraction sites, and datasets via grlc.io is available at cns-iu.github.io/hra-cell-type-populations-supporting-information#accessing-hrapop-data-via-hra-api.

### Visualizing CT populations for ASs, extraction sites, and datasets

A web interface to inspect HRApop Atlas CT populations via stacked bar graphs, entitled “HRApop Visualizer,” is available at apps.humanatlas.io/hra-pop-visualizer. A tutorial is provided at cns-iu.github.io/hra-cell-type-populations-supporting-information/#visualizing-cell-type-populations-for-as-es-and-datasets. An example screenshot is shown in **Fig. 6**.

**Fig. 6.**
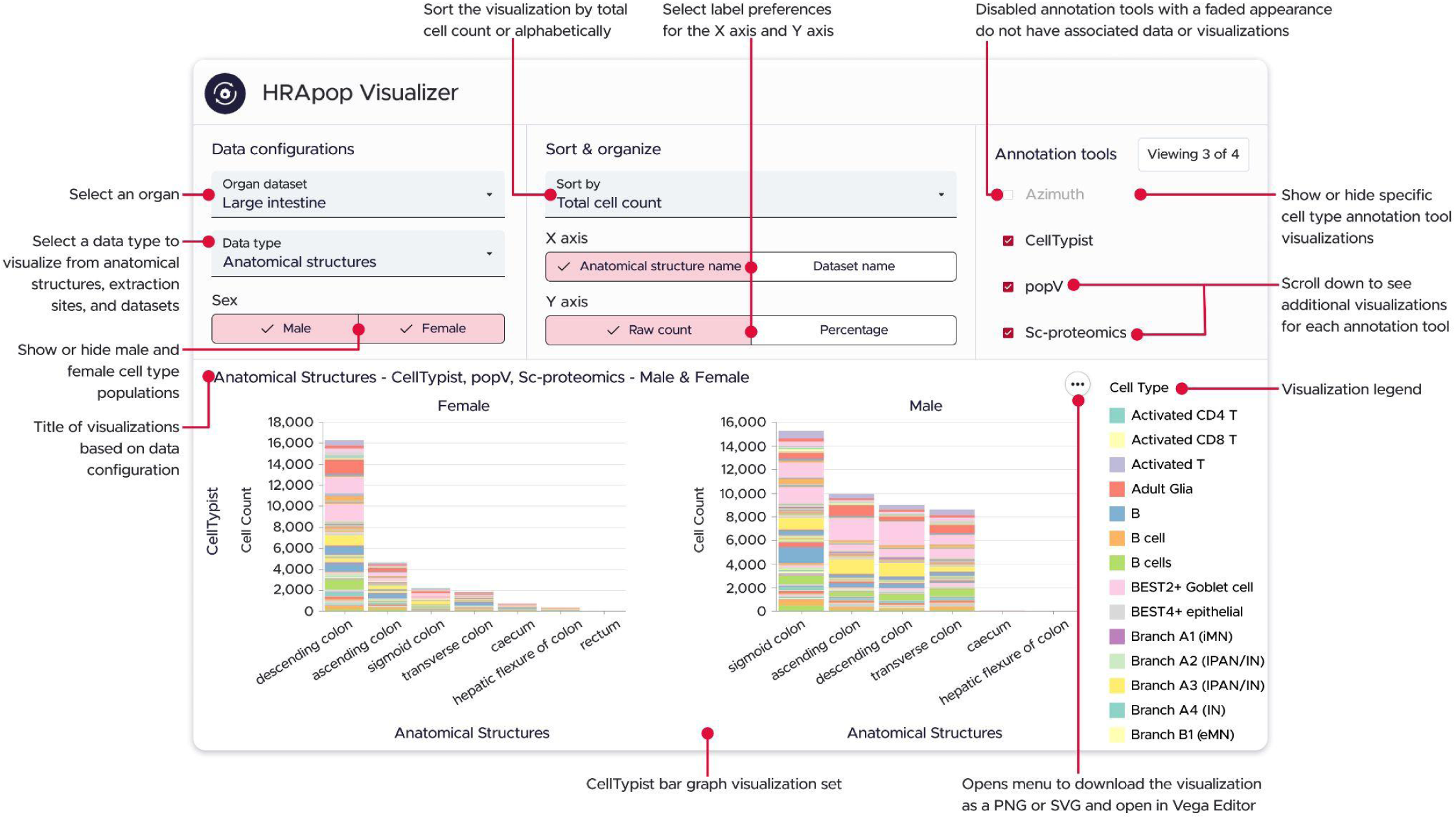
The HRApop Visualizer represents CT populations for ASs, extraction sites, and datasets as stacked bar graphs. For spatial reasons, only the CT populations from CellTypist are shown (those from popV and sc-proteomics data are not). Note that the rectum (left) and caecum/hepatic flexure of colon (right) do have bars, but the counts are so small that they are not rendered. Switching the y-axis to percentage makes these small counts visible as it changes the stacked bar graphs to 100% stacked bar graphs.

## Data Availability

Six major HRApop data products are available for download on Zenodo^82^. ASpop, DESpop, and corridors are also available via the HRA KG at purl.humanatlas.io/graph/hra-pop/v1.0 and on GitHub^123^, as well as in the form of canned SPARQL queries at apps.humanatlas.io/api/grlc/hra-pop.html. Links to all data are provided in **Table S1,** as is a complete listing of all code to construct and use HRApop. Examples for usage of this HRApop Atlas Data are available on the companion website at cns-iu.github.io/hra-cell-type-populations-supporting-information#usage-examples and the **Usage Notes** section.

## Code Availability

An overview of all GitHub repositories used to construct and use HRApop is provided in **Table S1**, separated by Data, Code (Construction), Code (Support), Collision Detection and Corridors, and Coverage and Visualization. All HRA UIs that expose/use HRApop are listed in **Table S2**. A companion website for this paper is available at cns-iu.github.io/hra-cell-type-populations-supporting-information and hosted in a GitHub repository^137^.

The Sankey diagram from **Fig. 2** can be explored interactively at cns-iu.github.io/hra-cell-type-populations-supporting-information/sankey_universe_plotly.html. A version showing only HRApop Atlas Data is shown in **Fig. S2**, with an interactive version available at cns-iu.github.io/hra-cell-type-populations-supporting-information/sankey_atlas_plotly.html. A companion website for this paper is available cns-iu.github.io/hra-cell-type-populations-supporting-information.

## Author Contributions

A.B. led the HRApop effort and the writing of this paper, compiled the figures, and provided input on the development of the DCTA Workflow. K.B. led the HRApop prototype development, led HRApop validation studies with support by M.G. and A.B., and co-wrote the manuscript. A.B. and B.W.H. implemented the RUI2CTpop Workflow. B.W.H. led the technical development of the DCTA and RUI2CT Workflows. L.C. implemented mesh-based collision detection and corridor generation. D.B. implemented the DCTA Workflow. D.Q. worked closely with tissue providers to register tissue blocks with the RUI, ran expert interviews, and developed and wrote up use cases. Y.J. provided guidance on sc-proteomics datasets as a generalized use case for HRApop. A.P.B. maintained and updated crosswalks from CT labels from CTann tools and author-provided CT labels for sc-proteomics datasets to CL plus higher-level CT aggregations. K.A. provided expert guidance on GTEx data analysis and usage. F.W. consulted on mesh-based collision detection and corridor generation.

## Competing Interests

The authors declare no competing interests.

## Acknowledgements

The authors would like to acknowledge work by Fauzan Isnaini, Aashay Gondalia, Vikrant Deshpande, Amber Ramesh, Nikhil Mahadevaswamy, and Mahadevan Narayanan Iyer on the HRApop prototype; Ellen M. Quardokus for authoring earlier CTann crosswalks; Vicky Daiya and Humaid Ilyas for their contributions to the DCTA Workflow and initial bar graph visualizations; Keyur Parekh and Juhi Khare for designing and implementing the HRApop Visualizer with input from Gauri Markandey; Devin Wright and Siddharth Apte for work on RUI registrations and extraction sites; Heidi Schlehlein and Kristen Browne for designing 3D reference objects; Ushma Patel, Tracey L. Theriault, Libby Maier, and Juhi Khare for figure design; and Niteesha Jangam for helping compile funding information.

The authors would also like to thank Yun (Renee) Zhang, Beverly Peng, Trang Nyuen, Thanh Long Nguyen, Richard S. Scheuermann, Nancy Ruschman, Humaid Ilyas, Supriya Bidanta, Elizabeth Ginexi, and Penny Cuda for their expert feedback on the near-final version of this paper, as well as Ellen M. Quardokus and Matthew R. Ruffalo for their feedback on earlier versions. They would also like to acknowledge expert input on CTann tools and crosswalks by Gesmira Molla, Skylar Li, Chuan Xu, and Can Ergen.

## Funding

The HRA is under active development by HuBMAP, SenNet, KPMP, GenitoUrinary Developmental Molecular Anatomy Project (GUDMAP)^138^, and the National Institute of Diabetes and Digestive and Kidney Diseases (NIDDK) with expert input by the HRA Editorial Board and in close collaboration with experts from 20+ other consortia. K.B. is a co-director of and is funded by the CIFAR MacMillan Multiscale Human program. K.B. is also supported via a Stiftung Charité Visiting Fellowship via Berlin Institute of Health at Charité (BIH).

This research has been supported by the following awards:

- The NIH Common Fund through the Office of Strategic Coordination/Office of the NIH Director:
  - HuBMAP:
    ▪ OT2OD026671 and OT2OD033756 (A.B., B.W.H., L.C., D.B., D.Q., M.G., Y.J., A.P.B., F.W., K.B.)
    ▪ OT2OD026675 (A.B., L.C. as NIH JumpStart Award)
    ▪ OT2OD033759 (A.B. as NIH JumpStart Fellowship)
  - SenNet: U24CA268108 (A.B., B.W.H., D.B., D.Q., M.G., Y.J., K.B.)
  - CFDE: OT2OD030545 (A.B., B.W.H., D.B., K.B.)
  - CFDE: 1R03OD039970-01 (A.B., B.W.H., D.B., M.G.)
- NIDDK:
  - KPMP: U01DK133090 (A.B., B.W.H., D.B., D.Q., K.B.)
  - U2CDK114886 (A.B., B.W.H., Y.J., K.B.)
  - U24DK135157 (B.W.H., K.B.);
- National Human Genome Research Institute (NHGRI):
  - GTEx: U24HG012090 (K.A.) ● National Cancer Institute (NCI):
  - U01CA242936 (L.C., F.W.)

This research uses IU’s computing infrastructure which was supported in part by Lilly Endowment, Inc., through its support for the Indiana University Pervasive Technology Institute.

The funders had no role in study design, data collection and analysis, decision to publish, or preparation of the manuscript. The content is solely the responsibility of the authors and does not necessarily represent the official views of the National Institutes of Health.

## Notes

https://cns-iu.github.io/hra-cell-type-populations-supporting-information

https://doi.org/10.5281/zenodo.17368990

https://doi.org/10.5281/zenodo.17368954

https://doi.org/10.5281/zenodo.17407573

https://app.globus.org/file-manager?origin_id=af603d86-eab9-4eec-bb1d-9d26556741bb&origin_path=/f53d60b5994333777a446dd7ad3b0304/extras/

https://doi.org/10.5281/zenodo.15603820

### Competing Interest Statement

The authors have declared no competing interest.

### Summary of Updates

For this revision, we restructured the paper, added new figures, and updated how we report HRApop numbers. It also contains more information on what data was selected and how it was validated.

