## Supplementary Information for "Cell type populations for 3D anatomical structures of the Human Reference Atlas"

### Supplementary Figures

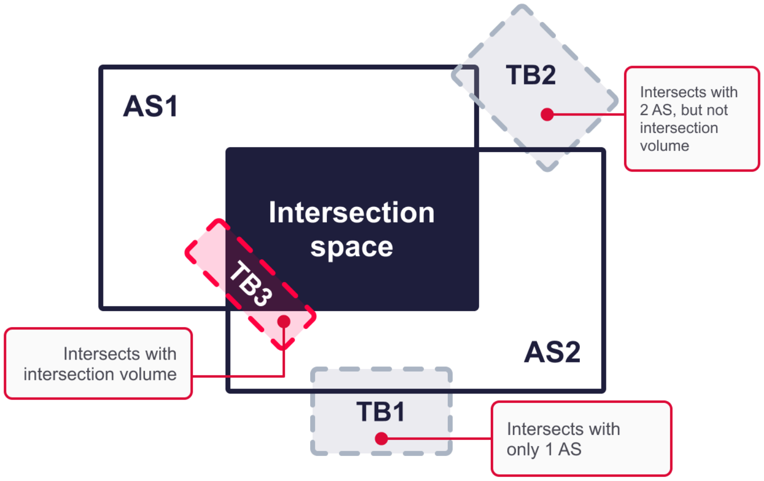


**Fig. S1**. **Intersection space and intersection volumes.** Two intersecting ASs create a 3D intersection space. Three exemplary tissue blocks intersect with one AS (TB1), both ASs (TB2), or both and their intersection space (TB3).

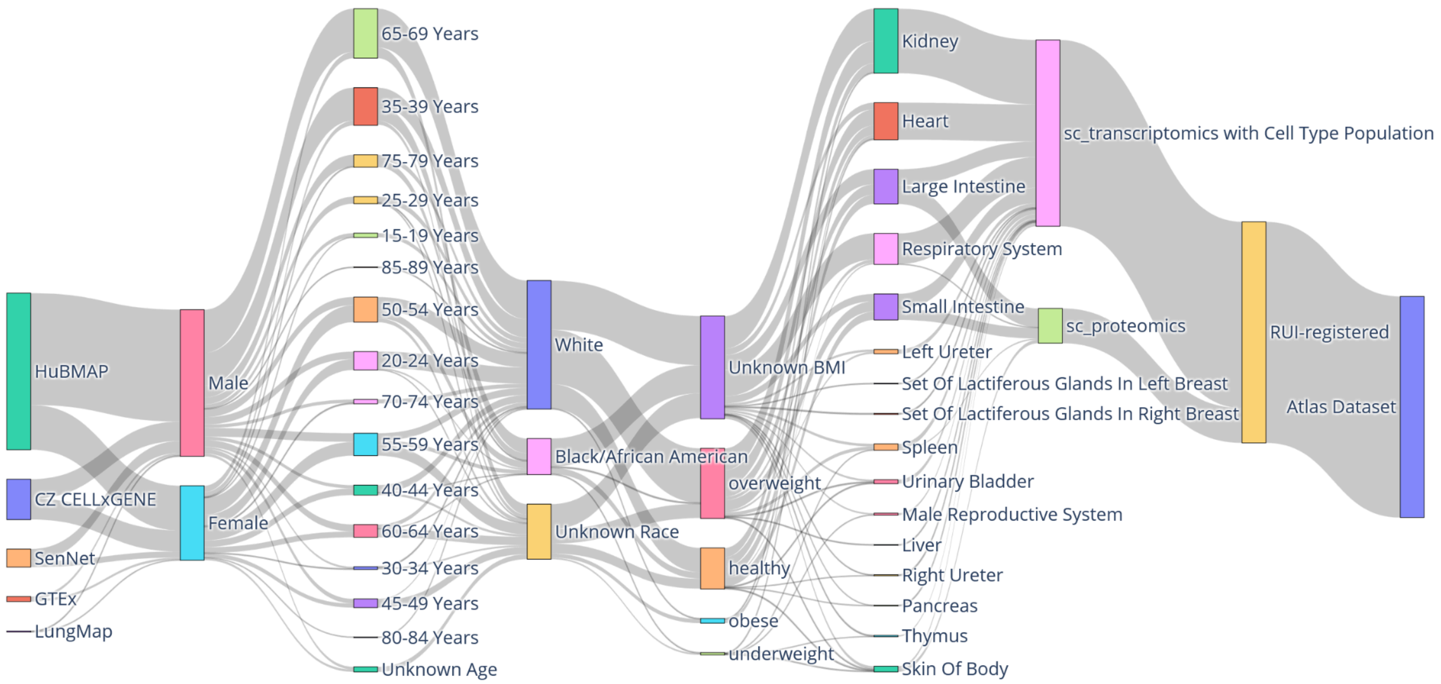


**Fig. S2. A Sankey Diagram for HRApop Atlas Data.** Interactive data visualization can be explored at cns-iu.github.io/hra-cell-type-populations-supporting-information/sankey_atlas_plotly.html.

### Supplementary Tables

**Table S1. Listing of all GitHub repositories used to construct and use HRApop.**

| Name | Description | URL |
|---|---|---|
| **Major Data Products** | | |
| DESpop | Provided in a variety of formats (including JSON-LD and Terse RDF Triple Language [Turtle][1]) via the HRA KG and as JSON-LD on GitHub | Zenodo[2]<br><br>GitHub (JSON-LD):<br>github.com/x-atlas-consortia/hra-pop/blob/main/output-data/v1.0/atlas-enriched-dataset-graph.jsonld<br><br>For datasets on GitHub (CSV):<br>github.com/x-atlas-consortia/hra-pop/blob/main/output-data/v1.0/reports/atlas-ad-hoc/cell-types-per-dataset.csv<br><br>For datasets on grlc.io (canned SPARQL query):<br>apps.humanatlas.io/api/grlc/hra-pop.html#get-/cell-types-per-dataset<br><br>For extraction sites on GitHub (CSV):<br>github.com/x-atlas-consortia/hra-pop/blob/main/output-data/v1.0/reports/atlas-ad-hoc/cell-types-per-extraction-site.csv<br><br>For extraction sites on grlc.io (canned SPARQL query):<br>apps.humanatlas.io/api/grlc/hra-pop.html#get-/cell-types-per-extraction-site<br><br>HRA KG:<br>cdn.humanatlas.io/digital-objects/graph/hra-pop/latest/assets/atlas-enriched-dataset-graph.jsonld |
| ASpop | Provided via the HRA KG and via GitHub | Zenodo[2]<br><br>HRA KG:<br>cdn.humanatlas.io/digital-objects/graph/hra-pop/v1.0/assets/atlas-as-cell-summaries.jsonld<br><br>GitHub (JSON-LD):<br>github.com/x-atlas-consortia/hra-pop/blob/main/output-data/v1.0/atlas-as-cell-summaries.jsonld<br><br>GitHub (CSV):<br>github.com/x-atlas-consortia/hra-pop/blob/main/output-data/v1.0/reports/atlas-ad-hoc/cell-types-in-anatomical-structurescts-per-as.csv<br><br>grlc.io (canned SPARQL query):<br>apps.humanatlas.io/api/grlc/hra-pop.html#get-/cell_types_in_anatomical_structurescts_per_as |
| Cell instances | Cell instances for sc-transcriptomics datasets in HRApop v1.0 | Zenodo[2] |

| | | |
|---|---|---|
| Cell instances with top-10,000 biomarkers | Cell instances for sc-transcriptomics datasets in HRApop v1.0 with top-10,000 biomarkers | Zenodo[2] |
| Corridors | Corridors for extraction sites for HRApop v1.0 | Zenodo[2]<br><br>GitHub: github.com/x-atlas-consortia/hra-pop/blob/main/output-data/v1.0/atlas-as-cell-summaries.jsonld<br><br>HRA KG: lod.humanatlas.io/graph/hra-pop/v1.0/ |
| HRApop quality control | ZIP folder with QC metrics for all 558 sc-transcriptomics datasets used in the DCTA Workflow | Zenodo[2] |
| **Code (Construction)** | | |
| Input data for DCTA and RUI2CTpop Workflows | Contains H5AD files (DCTA) and cell type populations for datasets + metadata (RUI2CTpop) | Globus[3] |
| DCTA Workflow | A set of scripts to download H5AD files, execute the Docker containers in HRApop CTann Tool Containers, and output CT populations and data metadata as input for the RUI2CTpop Workflow. | Release for HRApop v1.0: Zenodo[4]<br>Release for HRApop v1.0: GitHub[5]<br>Active repository: GitHub[6] |
| HRApop CTann Tool Containers | Has Docker containers for running CTann tools over H5AD files. | Release for HRApop v1.0: Zenodo[7]<br>Release for HRApop v1.0: GitHub[8]<br>Active repository: GitHub[9] |
| RUI2CTpop Workflow | A collection of scripts to compile HRApop from the output of the DCTA Workflow, which first provides CT populations and dataset metadata, then copies those files over to GitHub[10,11]. Scripts running over these input files are also on GitHub[12]. Output data is provided at github.com/x-atlas-consortia/hra-pop/tree/main/output-data/v1.0. For downstream analysis, reports based on | Release for HRApop v1.0: Zenodo[14]<br>Release for HRApop v1.0: GitHub[15]<br>Active repository: GitHub[11] |

| | | |
|---|---|---|
| | SPARQL queries are on GitHub[13].<br><br>A full log is available at raw.githubusercontent.com/x-atlas-consortia/hra-pop/refs/heads/main/output-data/v1.0/log.txt.<br><br>A file to capture the creation date of the pipeline having finished is at github.com/x-atlas-consortia/hra-pop/blob/main/output-data/v1.0/CREATION_DATE.<br><br>A Readme for the most recent run is present at github.com/x-atlas-consortia/hra-pop/blob/main/output-data/v1.0/README.md. | |
| Dataset info | Dataset IDs for all 16,293 datasets originally downloaded, incl. donor sex, assay type, CTann tool run, and unique CTs identified | GitHub[16] |
| CTann crosswalks | A collection of CSV files that link CT labels from Azimuth, CellTypist, and popV to CL or PCL IDs so they can be connected to the ASCT+B tables and other HRA Digital Objects. For example, the most recent crosswalk CSV file for Azimuth is available for download at the bottom of the HRA Digital Object landing page at lod.humanatlas.io/ctann/azimuth/latest. | lod.humanatlas.io/ctann |
| CTs level mapping | Report and queries to map crosswalked, unique CTs to higher-level CTs | Report:<br>github.com/x-atlas-consortia/hra-pop/blob/main/output-data/v1.0/reports/atlas-ad-hoc/cell-types-level-mapping.csv<br><br>Query:<br>github.com/x-atlas-consortia/hra-pop/blob/main/queries/atlas-ad-hoc/cell-types-level-mapping.rq |

| | | |
|---|---|---|
| | | Report (long): github.com/x-atlas-consortia/hra-pop/blob/main/output-data/v1.0/reports/atlas-ad-hoc/cell-types-level-mapping-long.csv<br><br>Query: github.com/x-atlas-consortia/hra-pop/blob/main/queries/atlas-ad-hoc/cell-types-level-mapping-long.rq |
| Not crosswalked | Reports for cell IDs that were not crosswalked during the DCTA Workflow | Input for RUI2CTpop Workflow: GitHub[17]<br><br>HRApop Atlas: GitHub[18] |
| CT populations and metadata for sc-transcriptomics and sc-proteomics data, produced by the DCTA Workflow and serving as input for the RUI2CTpop Workflow | **sc-transcriptomics-cell-summaries.jsonld.gz**: Contains CT populations for all sc-transcriptomics datasets annotated with a CTann tool in JSON-LD.<br><br>**sc-transcriptomics-dataset-metadata.csv**: Contains organ, donor metadata, . It also features a handler ID to denote which portal a dataset was downloaded from, a UUID, an assay type, and tissue provider information.<br><br>**sc-proteomics-cell-summaries.jsonld:** Contains CT populations for all sc-proteomics datasets in JSON-LD. The summaries are produced via the DCTA Workflow using the code at github.com/cns-iu/hra-node-dist-vis/blob/main/scripts/build-cell-summaries.js, which queries a listing of all possible sc-proteomics datasets (see GitHub[19]).<br><br>**sc-transcriptomics-dataset-metadata.csv:** Contains donor, tissue block, and data IDs for sc-proteomics datasets. | GitHub (all)[10]<br><br>GitHub (cell summaries for sc-protemics only)[20] |
| **Code (Support)** | | |
| HRA Registrations | Manually curated HRA Dataset Graphs. This repository holds static | GitHub[21] |

| | | |
|---|---|---|
| | dataset graphs for use in the EUI and other HRA applications. All registration Digital Objects are published to hubmapconsortium.github.io/hra-registrations/. All registrations are at hubmapconsortium.github.io/hra-registrations/**name-in-root-folder**/rui_locations.jsonld | |
| HRA Registrations Processor | Command line interface to simplify creating rui_locations.jsonld files | github.com/hubmapconsortium/hra-rui-locations-processor |
| Querying HRApop | A collection of SPARQL queries for use in this paper and for downstream analysis (to identify counts and get CT populations, biomarkers) | grlc.io/api-git/hubmapconsortium/ccf-grlc/subdir/hra-pop/ |
| RUI | Stand-alone RUI use to register tissue blocks | Stand-alone RUI: apps.humanatlas.io/rui/<br><br>RUI code repository: github.com/hubmapconsortium/hra-ui/tree/main/apps/ccf-rui |
| EUI | Deployed stand-alone RUI extraction site sets displayed in the EUI | hubmapconsortium.github.io/hra-registrations |
| CTann tools | Azimuth: v0.4.6<br>CellTypist: v1.6<br>popV: github.com/YosefLab/popV/tree/2d29c9a290d2015ec65ef0ef9f0e6b6d2277e7bb | Azimuth: azimuth.hubmapconsortium.org<br>CellTypist: www.celltypist.org<br>popV: github.com/YosefLab/PopV |
| **Collision Detection and Corridors** | | |
| HRA Mesh Collision API | Given an extraction site, get mesh-based collisions with the 3D reference object. | Code: github.com/hubmapconsortium/hra-tissue-block-annotation<br><br>Deployed: pfn8zf2gtu.us-east-2.awsapprunner.com/get-collisions<br><br>Public endpoint: apps.humanatlas.io/api/v1/collisions<br><br>Documentation for endpoint: apps.humanatlas.io/api/#post-/v1/collisions<br><br>Example result: github.com/hubmapconsortium/hra-tissue-block-annotation/blob/main/examples/test-registration-collisions.json |

| | | |
|---|---|---|
| 3D Corridor Generation API | Given an extraction site, generate a corridor with the 3D reference object as a GLB file | Code: github.com/hubmapconsortium/hra-corridor-generation<br><br>Deployed: dwwcpwad72.us-east-2.awsapprunner.com/get-corridor<br><br>Public endpoint: apps.humanatlas.io/api/v1/corridor<br><br>Documentation for endpoint: apps.humanatlas.io/api/#post-/v1/corridor |
| Corridors for downloading | Contains all corridor GLB files | github.com/x-atlas-consortia/hra-pop/tree/main/output-data/v1.0/corridors |
| Mesh-mesh collision and annotation | | Code: github.com/hubmapconsortium/hra-glb-mesh-collisions<br><br>On PyPi: pypi.org/project/hra-glb-mesh-collisions/0.1.0/ |
| **Coverage and Visualization** | | |
| CT Populations by AS | Provides a query for an overview of all AS-CT combinations in ASpop of HRApop v1.0, with sex, tool, CT, and cell percentage | apps.humanatlas.io/api/grlc/hra-pop.html#get-/cell_types_in_anatomical_structurescts_per_as. |
| Supporting Information for this paper | Contains assets for the companion website at cns-iu.github.io/hra-cell-type-populations-supporting-information/ | github.com/cns-iu/hra-cell-type-populations-supporting-information |
| Sankey Diagram for HRApop Atlas | Contains the deployed Sankey diagram for HRApop Atlas | cns-iu.github.io/hra-cell-type-populations-supporting-information/sankey_atlas_plotly.html |
| Sankey Diagram for input for RUI2CTpop Workflow | Contains the deployed Sankey diagram for the input for RUI2CTpop Workflow | cns-iu.github.io/hra-cell-type-populations-supporting-information/sankey_universe_plotly.html |
| HRApop Counts | Contains numbers reported in this paper | github.com/cns-iu/hra-cell-type-populations-supporting-information/blob/main/counts/hra_pop_counts.ipynb |

**Table S2. Listing of all HRA applications that use HRApop data.**

| Name | Description | URL[22] |
|---|---|---|
| FTU Explorer | Web-deployed UI to view and explore cell type populations from experimental datasets for 2D FTU illustrations | apps.humanatlas.io/ftu-explorer/#/ |
| HRA Organ Gallery[23,24] | Enables an immersive view of the HRA by showing 71 reference organs and 1,100+ tissue blocks alongside CT populations in VR[23,24]. | humanatlas.io/hra-organ-gallery |
| HRA API | Provides programmatic access to the HRA, enabling integration with other applications and services. A SPARQL endpoint for the HRA API allows users to write their own SPARQL queries. Canned queries for HRApop are available at apps.humanatlas.io/api/grlc/hra-pop.html. | API documentation: apps.humanatlas.io/api/<br><br>The HRA API on the HRA Portal: humanatlas.io/api |
| HRApop Visualizer | Web-deployed UI to visualize CT populations for 73 ASs, 230 extraction sites, and 662 HRApop Atlas datasets using stacked bar graphs, faceted by sex and CTann tool | https://apps.humanatlas.io/hra-pop-visualizer |

**Table S3. CTann settings.** Docker containers with full contexts for the three CTann tools are available on GitHub[25].

| Tool | Version | Code base | Models | Requirements |
|---|---|---|---|---|
| Azimuth | v0.4.6 | R | kidneyref/Kidney_L3/annotation.l3<br>lungref/Lung_v2_finest_level/ann_finest_level<br>heartref/Heart_L2/celltype.l2<br>humancortexref/subclass<br>pancreasref/Pancreas_L1/annotation.l1<br>pbmcref/Human_PBMC_L2/celltype.l2<br>bonemarrowref/Bone_marrow_L2/celltype.l2<br>adiposeref/Adipose_L2/celltype.l2 | anndata==0.9.1<br>contourpy==1.0.7<br>cycler==0.11.0<br>fonttools==4.39.4<br>h5py==3.8.0<br>importlib-metadata==6.6.0<br>importlib-resources==5.12.0<br>joblib==1.2.0<br>kiwisolver==1.4.4<br>llvmlite==0.40.1rc1<br>matplotlib==3.7.1<br>natsort==8.3.1<br>networkx==3.1<br>numba==0.57.0<br>numpy==1.24.3<br>packaging==23.1<br>pandas==2.0.2<br>patsy==0.5.3<br>Pillow==9.5.0<br>pynndescent==0.5.10<br>pyparsing==3.0.9<br>python-dateutil==2.8.2<br>pytz==2023.3<br>scanpy==1.9.3<br>scikit-learn==1.2.2<br>scipy==1.10.1<br>seaborn==0.12.2<br>session-info==1.0.0<br>six==1.16.0<br>statsmodels==0.14.0<br>stdlib-list==0.8.0<br>threadpoolctl==3.1.0<br>tqdm==4.65.0<br>tzdata==2023.3<br>umap-learn==0.5.3<br>zipp==3.15.0 |
| CellTypist | v1.6 | Python | Human_Lung_Atlas.pkl<br>Alternative: Cells_Lung_Airway.pkl<br>Adult_Human_Skin.pkl<br>Adult_Human_PancreaticIslet.pkl<br>Adult_Human_PancreaticIslet.pkl<br>Healthy_Adult_Heart.pkl<br>Healthy_Human_Liver.pkl<br>Human_AdultAged_Hippocampus.pkl<br>Human_Longitudinal_Hippocampus.pkl<br>Cells_Intestinal_Tract.pkl | anndata==0.9.2<br>celltypist==1.6.1<br>certifi==2023.7.22<br>charset-normalizer==3.3.1<br>click==8.1.7<br>contourpy==1.1.1<br>cycler==0.12.1<br>et-xmlfile==1.1.0<br>fonttools==4.43.1<br>h5py==3.10.0<br>idna==3.4<br>igraph==0.10.8<br>joblib==1.3.2<br>kiwisolver==1.4.5<br>leidenalg==0.10.1<br>llvmlite==0.41.1<br>matplotlib==3.8.0<br>natsort==8.4.0 |

| | | | | |
|---|---|---|---|---|
| | | | | networkx==3.2<br>numba==0.58.1<br>numpy==1.24.4<br>openpyxl==3.1.2<br>packaging==23.2<br>pandas==2.0.3<br>patsy==0.5.3<br>Pillow==10.1.0<br>pynndescent==0.5.10<br>pyparsing==3.1.1<br>python-dateutil==2.8.2<br>pytz==2023.3.post1<br>requests==2.31.0<br>scanpy==1.9.5<br>scikit-learn==1.3.2<br>scipy==1.11.3<br>seaborn==0.13.0<br>session-info==1.0.0<br>six==1.16.0<br>statsmodels==0.14.0<br>stdlib-list==0.9.0<br>texttable==1.7.0<br>threadpoolctl==3.2.0<br>tqdm==4.66.1<br>tzdata==2023.3<br>umap-learn==0.5.4<br>urllib3==2.0.7 |
| popV | See commit [github.com/YosefLab/popV/tree/2d29c9a290d2015ec65ef0ef9f0e6b6d2277e7bb](https://github.com/YosefLab/popV/tree/2d29c9a290d2015ec65ef0ef9f0e6b6d2277e7bb) | Python | model: Bladder<br>organ_level:urinary bladder<br><br>model: Blood<br>organ_level: blood<br><br>model: Bone_Marrow<br>organ_level: bone marrow<br><br>model: Eye<br>organ_level: eye<br><br>model: Fat<br>organ_level: adipose tissue<br><br>model: Large_Intestine<br>organ_level: large intestine<br><br>model: Liver<br>organ_level: liver<br><br>model: Lung<br>organ_level: lung<br><br>model: Lymph_Node<br>organ_level: mesenteric lymph node<br><br>model: Mammary<br>organ_level: mammary gland<br><br>model: Pancreas<br>organ_level: pancreas | absl-py==2.1.0<br>aiohappyeyeballs==2.4.0<br>aiohttp==3.10.5<br>aiosignal==1.3.1<br>anndata==0.10.9<br>annoy==1.17.3<br>array_api_compat==1.8<br>astunparse==1.6.3<br>attrs==24.2.0<br>bbknn==1.6.0<br>beautifulsoup4==4.12.3<br>celltypist==1.6.3<br>certifi==2024.8.30<br>charset-normalizer==3.3.2<br>chex==0.1.86<br>click==8.1.7<br>contextlib2==21.6.0<br>contourpy==1.3.0<br>cycler==0.12.1<br>Cython==3.0.11<br>docrep==0.3.2<br>et-xmlfile==1.1.0<br>etils==1.9.4<br>fbpca==1.0<br>filelock==3.15.4<br>flatbuffers==24.3.25<br>flax==0.9.0<br>fonttools==4.53.1<br>frozenlist==1.4.1<br>fsspec==2024.9.0<br>gast==0.6.0<br>gdown==5.2.0 |

| | | | | |
|---|---|---|---|---|
| | | | model: Prostate<br>Organ_level: prostate gland<br><br>model: Salivary Gland<br><br>model: Skin<br>Organ_level: skin<br><br>model: Small_Intestine<br>organ_level: small intestine<br><br>model: Spleen<br>organ_level: spleen<br><br>model: Thymus<br>organ_level: thymus<br><br>model: Tongue<br><br>model: Trachea<br>organ_level: trachea<br><br>model: Uterus<br>organ_level: uterus<br><br>model: Vasculature<br>organ_level: blood vasculature | geosketch==1.2<br>google-pasta==0.2.0<br>grpcio==1.66.1<br>h5py==3.11.0<br>harmony-pytorch==0.1.8<br>huggingface-hub==0.24.6<br>humanize==4.10.0<br>idna==3.8<br>igraph==0.11.6<br>importlib_resources==6.4.4<br>intervaltree==3.1.0<br>jax==0.4.31<br>jaxlib==0.4.31<br>Jinja2==3.1.4<br>joblib==1.4.2<br>keras==3.5.0<br>kiwisolver==1.4.7<br>legacy-api-wrap==1.4<br>leidenalg==0.10.2<br>libclang==18.1.1<br>lightning==2.1.4<br>lightning-utilities==0.11.7<br>llvmlite==0.43.0<br>Markdown==3.7<br>markdown-it-py==3.0.0<br>MarkupSafe==2.1.5<br>matplotlib==3.9.2<br>mdurl==0.1.2<br>ml-dtypes==0.4.0<br>ml_collections==0.1.1<br>mpmath==1.3.0<br>msgpack==1.0.8<br>mudata==0.3.1<br>multidict==6.0.5<br>multipledispatch==1.0.0<br>namex==0.0.8<br>natsort==8.4.0<br>nest-asyncio==1.6.0<br>networkx==3.3<br>numba==0.60.0<br>numpy==1.26.4<br>numpyro==0.15.2<br>nvidia-cublas-cu12==12.1.3.1<br>nvidia-cuda-cupti-cu12==12.1.105<br>nvidia-cuda-nvrtc-cu12==12.1.105<br>nvidia-cuda-runtime-cu12==12.1.10<br>5<br>nvidia-cudnn-cu12==9.1.0.70<br>nvidia-cufft-cu12==11.0.2.54<br>nvidia-curand-cu12==10.3.2.106<br>nvidia-cusolver-cu12==11.4.5.107<br>nvidia-cusparse-cu12==12.1.0.106<br>nvidia-nccl-cu12==2.20.5<br>nvidia-nvjitlink-cu12==12.6.68<br>nvidia-nvtx-cu12==12.1.105<br>obonet==1.1.0<br>OnClass==1.3<br>openpyxl==3.1.5<br>opt-einsum==3.3.0<br>optax==0.2.3 |

| | | | | |
|---|---|---|---|---|
| | | | | optree==0.12.1<br>orbax-checkpoint==0.6.1<br>packaging==24.1<br>pandas==1.5.3<br>patsy==0.5.6<br>pillow==10.4.0<br>PopV @ git+github.com/czbiohub/PopV@2d29c9a290d2015ec65ef0ef9f0e6b6d2277e7bb<br>protobuf==4.25.4<br>psutil==6.0.0<br>Pygments==2.18.0<br>pynndescent==0.5.13<br>pyparsing==3.1.4<br>pyro-api==0.1.2<br>pyro-ppl==1.9.1<br>PySocks==1.7.1<br>python-dateutil==2.9.0.post0<br>pytorch-lightning==2.4.0<br>pytz==2024.1<br>PyYAML==6.0.2<br>regex==2024.7.24<br>requests==2.32.3<br>rich==13.8.0<br>safetensors==0.4.5<br>scanorama==1.7.4<br>scanpy==1.10.2<br>scikit-learn==1.1.3<br>scikit-misc==0.5.1<br>scipy==1.14.1<br>scvi-tools==1.1.6<br>seaborn==0.13.2<br>sentence-transformers==3.0.1<br>session_info==1.0.0<br>six==1.16.0<br>sortedcontainers==2.4.0<br>soupsieve==2.6<br>statsmodels==0.14.2<br>stdlib-list==0.10.0<br>sympy==1.13.2<br>tensorboard==2.17.1<br>tensorboard-data-server==0.7.2<br>tensorflow==2.17.0<br>tensorflow-io-gcs-filesystem==0.37.1<br>tensorstore==0.1.65<br>termcolor==2.4.0<br>texttable==1.7.0<br>threadpoolctl==3.5.0<br>tokenizers==0.19.1<br>toolz==0.12.1<br>torch==2.4.1<br>torchmetrics==1.4.1<br>tqdm==4.66.5<br>transformers==4.44.2<br>triton==3.0.0<br>typing_extensions==4.12.2<br>umap-learn==0.5.6<br>urllib3==2.2.2 |

| | | | | |
|---|---|---|---|---|
| | | | | Werkzeug==3.0.4<br>wget==3.2<br>wrapt==1.16.0<br>yarl==1.9.11<br>zenodo-get==1.5.1<br>zipp==3.20.1 |

**Table S4. Reports about dataset and cell counts from the RUI2CTpop Workflow.**

| **Counts** | | |
|---|---|---|
| Input for RUI2CTpop Workflow: sc-transcriptomics cell counts | Contains the number of annotated cells in sc-transcriptomics datasets that are part of the input for RUI2CTpop Workflow | github.com/x-atlas-consortia/hra-pop/blob/main/output-data/v1.0/reports/universe-ad-hoc/universe-sc-transcriptomics-cell-counts.csv |
| Input for RUI2CTpop Workflow: sc-transcriptomics cell counts (with pre-annotated cell counts) | Contains the number of annotated cells in sc-transcriptomics datasets that are part of the input for RUI2CTpop Workflow that have been mapped via CTann tools (see **Box 1**). Pre-annotated cell counts are the number of rows in the cell by gene matrix of the H5AD file. Some cells may not get annotated for various reasons, thus the discrepancy | github.com/x-atlas-consortia/hra-pop/blob/main/output-data/v1.0/reports/universe-ad-hoc/universe-sc-transcriptomics-cell-instance-counts.csv |
| Input for RUI2CTpop Workflow: sc-proteomics cell counts | Contains the number of annotated cells in sc-proteomics datasets that are part of the HRApop Atlas | github.com/x-atlas-consortia/hra-pop/blob/main/output-data/v1.0/reports/universe-ad-hoc/universe-sc-proteomics-cell-counts.csv |
| HRApop Atlas: sc-transcriptomics cell counts | Contains the number of annotated cells in sc-transcriptomics datasets that are part of the HRApop Atlas | github.com/x-atlas-consortia/hra-pop/blob/main/output-data/v1.0/reports/atlas-ad-hoc/atlas-sc-transcriptomics-cell-counts.csv |
| HRApop Atlas: sc-proteomics cell counts | Contains the number of annotated cells in sc-proteomics datasets that are part of the HRApop Atlas | github.com/x-atlas-consortia/hra-pop/blob/main/output-data/v1.0/reports/atlas-ad-hoc/atlas-sc-proteomics-cell-counts.csv |
| HRApop Atlas: Number of extraction sites per AS and intersection volume/percentage of the extraction site (see **Box 1**) | Lists extraction site, organ/AS, intersection volume, AS volume and tissue block volume. | github.com/x-atlas-consortia/hra-pop/blob/main/output-data/v1.0/reports/atlas/table-s5.csv |
| SPARQL queries | Contains SPARQL queries for all reports generated in in the output data folder on GitHub[26] | github.com/x-atlas-consortia/hra-pop/tree/main/queries |

**Table S5. CT choices.** 26 of the 201 CTs were classified under two different high-level CTs. This table shows the decision process used to select the most appropriate grouping class for each CT.

| CT | Chosen Category | Non-chosen category | Rationale |
|---|---|---|---|
| hematopoietic stem cell | hematopoietic cell | stem cell | Reflects lineage identity central to its biological role |
| neuroendocrine cell | epithelial cell | neural cell | Developmentally and structurally epithelial |
| pancreatic D cell | epithelial cell | neural cell | Developmentally and structurally epithelial |
| intestinal crypt stem cell of large intestine | stem cell | epithelial cell | Stemness prioritized as foundational to epithelial renewal |
| intestinal crypt stem cell of small intestine | stem cell | epithelial cell | Stemness prioritized as foundational to epithelial renewal |
| lung interstitial macrophage | hematopoietic cell | connective tissue cell | Lineage from hematopoietic system prioritized over location |
| pulmonary neuroendocrine cell | epithelial cell | neural cell | Developmentally and structurally epithelial |
| granulocyte monocyte progenitor cell | hematopoietic cell | bone cell | Reflects hematopoietic lineage over location in the bone marrow |
| mesenchymal stem cell of adipose tissue | stem cell | connective tissue cell | Stemness and multipotency prioritized over local connective tissue association. |
| mesenchymal stem cell of abdominal adipose tissue | stem cell | connective tissue cell | Stemness and multipotency prioritized over local connective tissue association. |
| central nervous system macrophage | hematopoietic cell | neural cell | Immune lineage prioritized over CNS location |
| vascular leptomeningeal cell | connective tissue cell | neural cell | Mesenchymal origin and structural connective function prioritized over CNS location |
| mature microglial cell | hematopoietic cell | neural cell | Immune lineage prioritized over CNS location |
| ependymal cell | neural cell | epithelial cell | Derived from neuroectoderm and part of CNS despite epithelial morphology |
| smooth muscle cell of the brain vasculature | muscular cell | neural cell | Reflects muscular identity, over location |
| mesenchymal stem cell | stem cell | connective tissue cell | Stemness and multipotency prioritized over local connective tissue association. |
| basal cell of epidermis | stem cell | epithelial cell | Stemness emphasized as central to epidermal renewal |

| | | | |
|---|---|---|---|
| choroid plexus epithelial cell | neural cell | epithelial cell | Derived from neuroectoderm and part of CNS despite epithelial morphology |
| microglial cell | hematopoietic cell | neural cell | Primitive macrophage origin prioritized over CNS-resident role |
| type D cell of colon | epithelial cell | neural cell | Developmentally and structurally epithelial |
| type EC enteroendocrine cell | epithelial cell | neural cell | Developmentally and structurally epithelial |
| cycling type EC enteroendocrine cell | epithelial cell | neural cell | Developmentally and structurally epithelial |
| type D cell of small intestine | epithelial cell | neural cell | Developmentally and structurally epithelial |
| retinal blood vessel endothelial cell | endothelial cell | neural cell | Endothelial identity and vascular role prioritized over retinal location |
| retinal pigment epithelial cell | epithelial cell | neural cell | Morphology and function as barrier epithelium prioritized over retinal location |
| endosteal cell | bone cell | epithelial cell | Skeletal lineage and role in bone homeostasis prioritized over lining morphology |
